# Adaptive responses to antifungal selection depend on dose and drug combination in experimentally evolved yeast

**DOI:** 10.64898/2026.10.28.759463

**Authors:** Megan Sandoval-Powers, Giovanni Crestani, Hannah Dugo, Ellery Shearman, Molly K. Burke

## Abstract

Natural populations rarely face a single stressor at a single intensity, yet most laboratory evolution experiments impose exactly that. Theory predicts that stronger or compounded selection pressures should change the magnitude of adaptive change as well as the specific loci involved, but this prediction is rarely tested directly. Here, we use an evolve-and-resequence approach to compare genomic responses in *Saccharomyces cerevisiae* across antifungal environments differing in the concentration and number of drugs applied. Twelve replicate populations founded from a genetically diverse ancestor were evolved for >50 generations under low and high concentrations of caspofungin or clotrimazole alone, a combination of the two drugs, or no drug, with pooled-population sequencing at three timepoints. We found little overlap among candidate loci between concentrations of the same drug or between single and combined treatments. Alleles favored in one environment did not shift predictably in the others, departing from expectations of additivity. We also observed that most candidate loci fell outside known antifungal resistance genes, except for *FKS1* in all caspofungin treatments and a putative gain of the *ERG11*-containing chromosome VIII in 9 of 12 populations evolved under high clotrimazole. Remaining candidates implicated diverse cellular functions that differed among environments. Furthermore, we found little evidence of fitness costs in drug-free medium across all evolved populations. Our results collectively show that stronger and combined antifungal selection redirects genomic responses rather than simply amplifying those favored under less challenging conditions, highlighting the limits of predicting adaptation from simpler selection regimes.

**Significance statement:** Predicting evolutionary responses across environmental contexts remains a major challenge. Populations exposed to similar stresses often show repeatable patterns of adaptation, but whether these patterns persist as stress intensifies or multiple stresses occur together is less clear. By evolving yeast under different concentrations of two antifungal drugs, alone and in combination, we found that the genetic changes responding to selection differed substantially across treatments. Stronger and combined antifungal selection favored distinct genomic responses beyond the drugs’ primary targets, while retaining some shared responses at known resistance-associated loci. These findings show that evolutionary responses observed under one set of conditions may be poor predictors of adaptation under stronger or more complex stress, with implications for anticipating antimicrobial resistance across different drug concentrations and combination therapies.

## Introduction

Ecological processes and anthropogenic change continually alter the abiotic and biotic conditions experienced by populations, creating dynamic selective landscapes that shape adaptation (Otto 2018; Pirotta et al. 2022; Rêgo et al. 2024). These landscapes vary not only through time and space, but also in the intensity and number of selective pressures they impose (Kingsolver et al. 2001; Anderson et al. 2014; Munson et al. 2025). Although theoretical models have explored how adaptation should proceed under diverse environmental conditions (Chevin et al. 2010; Jain and Stephan 2017; Stetter et al. 2018), empirical tests of these predictions are much more limited. We therefore know relatively little about how selection intensity and environmental complexity influence the magnitude and genetic basis of adaptation.

Understanding these effects is important for predicting responses to environmental change, with implications for conservation biology (Nadeau and Urban 2019; Brennan et al. 2022), the evolution of antimicrobial resistance (Schmidlin et al. 2024), and the maintenance of heritable variation (Sakamoto and Yeaman 2026).

Experimental evolution provides a powerful means to test questions about the nature of selection and evolutionary dynamics. In this approach, experimental populations are evolved across multiple generations under controlled conditions in which one or more environmental factors are systematically manipulated (Kawecki et al. 2012). This approach helps address a major challenge in natural systems, where multiple environmental factors change simultaneously and are difficult to isolate and control (Vandenkoornhuyse et al. 2010; Rêgo et al. 2024). When combined with whole-genome sequencing, experimental evolution can be used to track allele frequency changes over time and identify genomic regions responding to selection through an approach known as evolve-and-resequence (E&R) (Long et al. 2015; Schlötterer et al. 2015).

E&R studies have considerably advanced our understanding of adaptive genetic architecture, but most have focused on relatively simple environments, that rarely include treatments differing in strength or with interacting selective pressures (Bailey and Bataillon 2016; Graf et al. 2021).

This bias is in part due to the practical demands of these experiments, which require additional populations and sequencing effort for each treatment. Consequently, we still have limited experimental evidence of how well patterns of adaptation observed under simple conditions predict responses to stronger or more complex selection.

One key question is whether stronger selection amplifies an existing genomic response or changes which loci contribute to adaptation. Models of polygenic adaptation generally predict that as selection intensifies, the response shifts from subtle, coordinated allele frequency changes toward larger, more rapid shifts at selected loci (Jain and Stephan 2017; Barghi and Schlötterer 2020). Simulations seem to support this idea, showing that populations adapting toward more distant phenotypic optimum can undergo larger allele frequency changes at target loci, with some beneficial alleles reaching fixation (Stetter et al. 2018; Christodoulaki et al. 2019). These expectations often rely on additive genetic effects, under which stronger selection favors the same variants but drives them further in the favored direction. However, additive expectations may break down if alleles driving adaptation under strong selection become neutral or costly under weaker selection, as can occur through epistasis or context-dependent pleiotropy (Visscher and Yang 2016; Barton 2017). Under these circumstances, increasing selection intensity could change which variants are favored rather than simply strengthening an existing response.

Empirical evidence is limited, but in a recent study by Phillips et al. (2026), the authors found that high ethanol stress in yeast increased the magnitude of allele frequency change at loci associated with ethanol adaptation but also engaged partially distinct biological pathways compared to moderate ethanol stress. More indirect evidence comes from *Drosophila simulans*, where populations adapting to hot and cold temperature regimes shared almost no selection targets (Otte et al. 2021). Notably, these studies have primarily focused on non-specific stressors such as ethanol and temperature that affect many cellular processes at once (Wolf et al. 2023; Li et al. 2025). For stressors with more specific cellular targets, such as antimicrobial drugs, adaptive pathways may be more constrained, potentially leading to greater overlap in responding loci across stress intensities. The extent to which stronger selection amplifies existing responses or favors alternative genetic solutions remains an open question, particularly for targeted xenobiotic stressors.

Environments can also become more challenging when populations experience multiple selective pressures simultaneously, as adaptation may reflect not only the effects of individual stressors but also their interactions (Brennan et al. 2022; Pirotta et al. 2022). If stressors act additively, combined exposure might be expected to amplify allele frequency changes at loci favored under each stressor alone. Alternatively, multiple stressors may also interact synergistically or antagonistically, creating selective conditions distinct from either individual stressor (Brennan et al. 2022; Burny et al. 2022). Such non-additive interactions may shift the targets of selection depending on which alleles confer the greatest net benefit under the combined conditions. Although a limited number of E&R studies have begun to examine responses to combined stressors, we still lack a clear understanding of how multiple stressors reshape the genetic architecture of adaptation and the extent to which their effects can be predicted from single-stressor responses.

Antifungal drugs offer a practical and clinically relevant way for examining these questions because drug identity, concentration, and combination can be manipulated independently. Different antifungal classes act on distinct cellular targets (K Mazu et al. 2016), allowing selection intensity and complexity to be varied within the same experimental framework. These conditions are also relevant to fungal populations in clinical and natural environments, where drug exposure can vary and combination therapies are used to improve treatment outcomes and limit resistance (Chait et al. 2016; Todd et al. 2023; Bognár et al. 2024). *Saccharomyces cerevisiae* is well suited to this approach because many of its antifungal resistance mechanisms are well-characterized, and it shares important resistance genes with pathogenic fungi (Anderson et al. 2003; Marie and White 2009). This makes it possible to assess whether adaptation follows established resistance mechanisms or involves less expected genetic changes under different treatments. Moreover, outcrossed *S. cerevisiae* populations can be leveraged that harbor substantial standing genetic variation (Cubillos et al. 2009; Burke et al. 2020), allowing us to track selection across many segregating loci and examine adaption that does not depend solely on newly arising mutations.

Here, we used an E&R approach to examine genomic and phenotypic adaptation in outcrossed *S. cerevisiae* populations evolved under low or high concentrations of two antifungals, caspofungin acetate (CAS) and clotrimazole (CLO), both individually and in combination (CASCLO). CAS is an echinocandin that inhibits fungal cell wall synthesis by inhibition of the enzyme (1,3)-β-D-glucan synthase (Letscher-Bru and Herbrecht 2003), whereas CLO is an azole that disrupts ergosterol biosynthesis and affects the fungal cell membrane (Crowley and Gallagher 2014). Using these drugs allowed us to compare adaptation to stressors with different cellular targets with well-known modes of action. We tracked allele frequency changes over time to determine whether stronger selection and combined selection increased the magnitude of genomic responses at selected loci and whether the targets of selection differed across treatments. We also compared candidate loci with known antifungal resistance genes to assess whether stronger or combined selection was more likely to engage previously characterized resistance targets. Finally, we measured the growth of evolved populations across drug and benign environments to evaluate changes in drug tolerance and potential fitness trade-offs. We found that increasing drug concentration or combining the two drugs did not predictably extend the responses observed under less challenging conditions but instead generated largely context-dependent genomic trajectories.

## Results

To compare how populations adapt to stronger and combined antifungal selection, we evolved 12 replicate populations of a genetically diverse, outcrossed *S. cerevisiae* ancestor under six selective regimes that differed in the strength and number of antifungal pressures applied (**Fig. 1**). Populations were exposed to low or high concentrations of caspofungin (Low CAS, High CAS) or clotrimazole (Low CLO, High CLO), or to the two low concentrations combined (CASCLO). Concentrations were determined from dose-response assays to impose distinct levels of stress while still permitting sufficient growth for serial transfer (**Fig. S1**). A no-drug DMSO control was included to account for background adaptation to laboratory conditions, yielding 72 evolved populations in total. Populations were sampled at three timepoints (T_1_, T_7_, T_14_), generating 216 samples (72 evolved populations × 3 timepoints) that were sequenced via pooled-population whole-genome sequencing, enabling inference of genome-wide allele frequency dynamics over time. T_14_ populations were additionally phenotyped to assess growth performance across treatments. Estimated cumulative generations ranged from 54–69 across treatments (**Table 1**), with variation arising primarily from differences in growth rates during early transfers. After filtering for read depth, ancestral polymorphism, and low-confidence sites based on founder genotypes, pooled sequencing yielded 64,764 high-quality biallelic SNPs at a median genome-wide coverage of 86× (range: 27×–290×; **Table S1**). This filtered dataset was used to identify loci responding to selection and track allele frequency dynamics across treatments.

**Figure 1.**
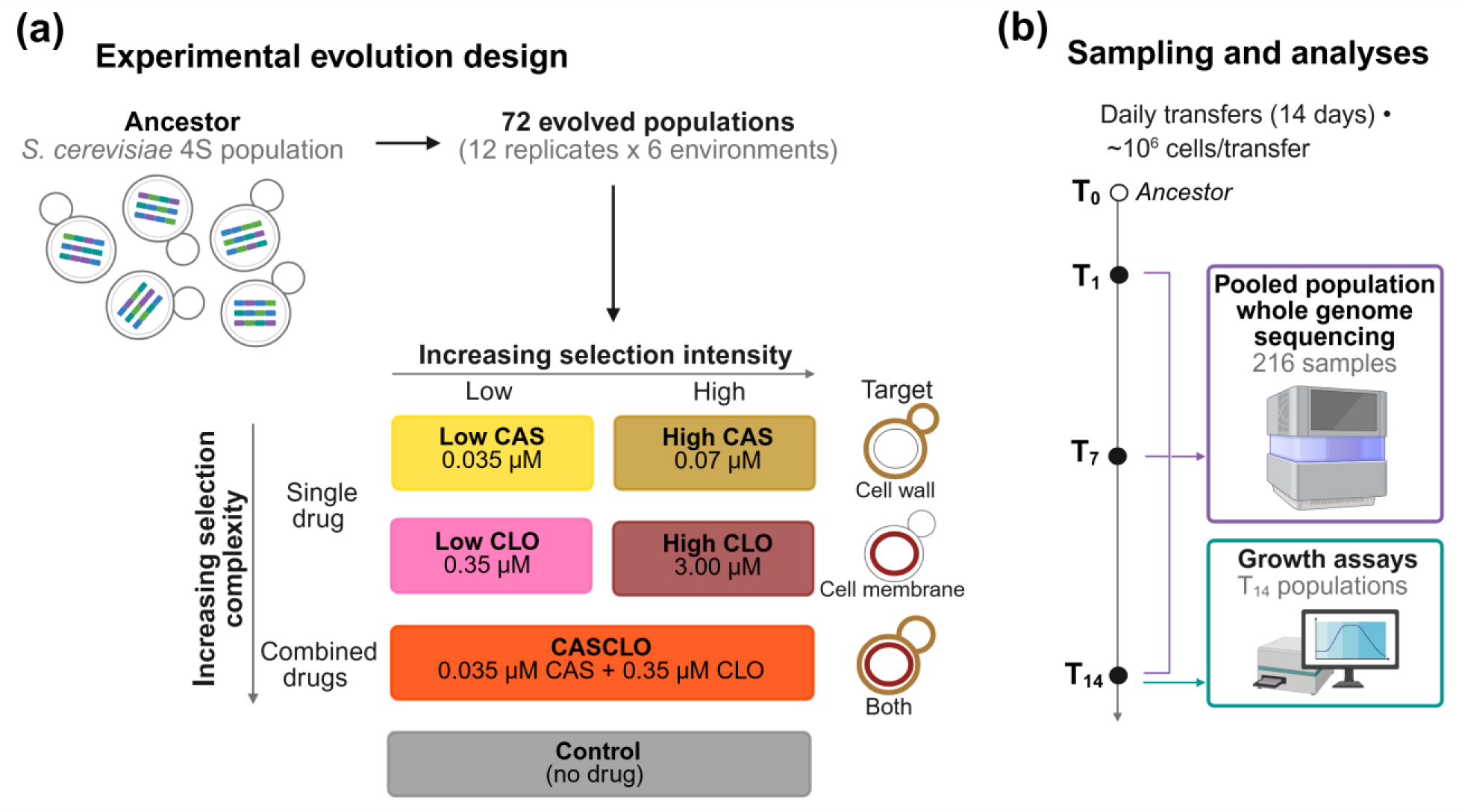
Experimental design. (a) An outcrossed *S. cerevisiae* population (4S) was used to found 72 populations, evolved under each of six environments: low and high concentrations of caspofungin (CAS; cell wall target) and clotrimazole (CLO; cell membrane target), a combination of the low concentrations of both drugs (CASCLO), and a no-drug Control. Environments varied in selection intensity (low vs. high concentration) and complexity (single vs. combined drugs). (b) Populations were propagated by daily transfer into fresh selective media of ∼10⁶ cells for 14 days. Pooled whole-genome sequencing was performed at T_1_, T_7_, and T_14_ (216 samples), with the ancestor serving as T_0_, and growth assays were performed on T_14_ populations. Created in BioRender. Sandoval-Powers, M. (2026) https://BioRender.com/buz5nqr

**Table 1.**
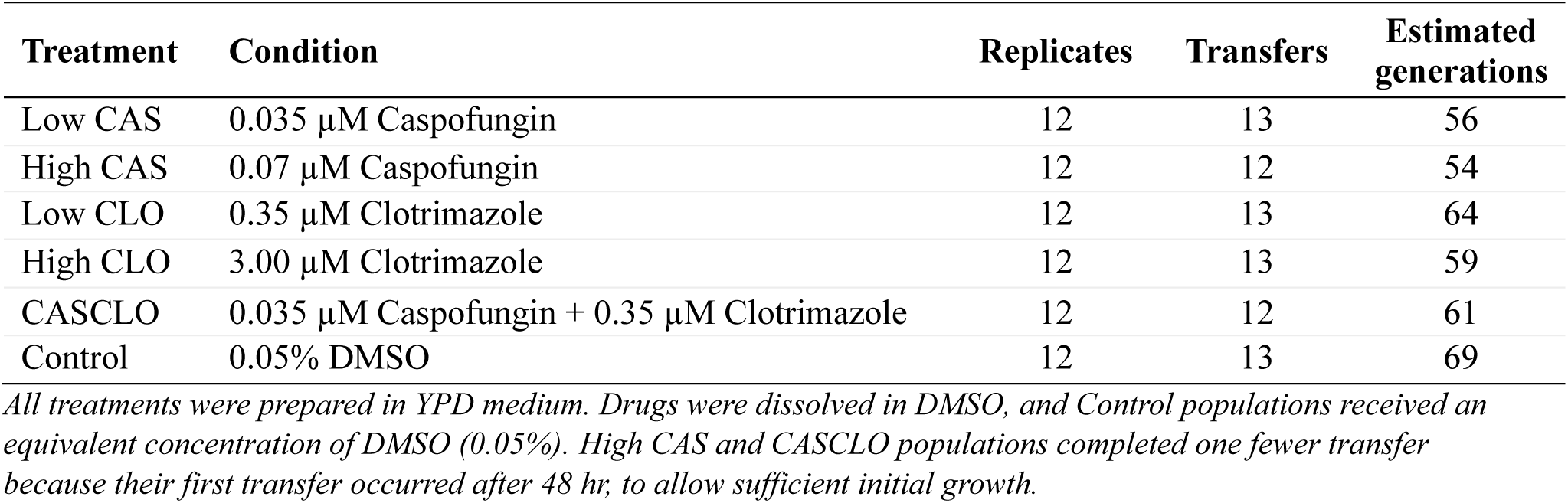
Summary of treatment conditions and estimated generations during experimental evolution.

### Related antifungal regimes produce distinct genome-wide responses, with early fixation concentrated under strong selection

To broadly characterize genome-wide responses across selection regimes, we compared allele frequency patterns among treatments and timepoints using principal component analysis (PCA) and site frequency spectra (SFS). Most drug-evolved populations progressively diverged from the ancestor and Controls. Divergence was greatest under High CAS and High CLO, where replicate populations were also more dispersed, indicating that stronger selection produced larger and more variable genome-wide responses (**Fig. S2**). Low CLO populations were an exception, remaining closely clustered with Controls and showing limited genome-wide differentiation (**Fig. S2b**).

Because Low CLO populations showed such a limited response, we chose to compare genome-wide differentiation between the combined drug populations (CASCLO) and those grown under the high single-drug treatments (High CAS and High CLO) to make more direct comparisons between the most stressful treatments (**Fig. S1**) and indirectly minimize biases due to differences driven simply by weaker selection. We use this comparison here and in several analyses that follow. By T_14_, CASCLO populations overlapped with High CLO along PC2 while separating from High CAS along the same axis, and showed considerably more variation along PC1 (**Fig. S2c**).

Site frequency spectra at T_14_ also showed differences in allele frequency distributions among treatments. Drug-evolved populations had more variants approaching loss or fixation than Controls (AF near 0 or 1; **Fig. S3**), except again for Low CLO populations. Across all treatments, however, many variants remained at intermediate frequencies, indicating that substantial standing genetic variation persisted throughout the experiment.

A small subset of variants showed an extreme form of allele frequency change, reaching early fixation or loss (allele frequency of 1 or 0) in every replicate of a given treatment by T_1_ and remaining fixed through T_14_ (hereafter early-fixed sites). These sites were rare. Most SNPs remained polymorphic at T_1_, and many persisted at intermediate frequencies through T_14_, indicating that standing variation was not broadly depleted early in the experiment. Because these sites reached extreme frequencies before the first sampled timepoint, their trajectories could not be estimated with the temporal models used for other loci. We nonetheless retained them as potential signals of selection, as several were found to occur in genes relevant to the drugs, including *FKS1*. To isolate drug-associated changes, we excluded sites that also fixed by T_1_ in Control populations, likely reflecting laboratory adaptation, and sites showing minimal change from the ancestor, which were already near fixation. Of the 64,764 SNPs, 755 (∼1.2%) met these criteria in at least one treatment. Early-fixed sites were most common under high drug treatments. CAS populations had 551 of these sites under High CAS and 140 under Low CAS, with 121 shared between concentrations. CLO populations showed far fewer, with 24 under High CLO and 15 under Low CLO, of which 5 were shared. CASCLO populations contained 507 early-fixed sites, of which 311 were shared with High CAS, 5 with High CLO, and 184 were specific to CASCLO.

### Responses at candidate loci do not scale predictably with stronger or combined selection

To test whether stronger or combined antifungal selection amplified responses observed under milder conditions, we modeled allele frequency change over time at each SNP using quasibinomial generalized linear models (GLMs), excluding the early-fixed sites described above. For each treatment, SNPs were ranked by FDR-adjusted *p*-values, and the top 5% representing approximately ∼3,130 SNPs per treatment were designated as candidate loci. We quantified allele frequency change using the GLM coefficient (β), which represents the rate of change on the logit scale. For each candidate, we identified the allele that increased in the treatment where it was detected and measured how that same allele changed in the other treatments. Responses were summarized as median effect sizes (β̃) with interquartile ranges (IQR) and compared among treatments using Friedman rank-sum tests with paired Wilcoxon contrasts (Holm-adjusted). Given the large sample size per treatment, even small differences in β̃ were statistically significant, so we interpret responses primarily by their magnitude.

Responses at CAS candidate loci did not increase with stronger CAS selection. Low CAS candidates shifted strongly under Low CAS (β̃ = 0.241, IQR: 0.167–0.320) but barely changed under High CAS (β̃ = 0.003, IQR -0.027–0.043, compared with -0.008 in Controls; **Fig. 2a**). High CAS candidates shifted strongly under High CAS (β̃ = 0.165, IQR 0.122–0.274) and more modestly under Low CAS (β̃ = 0.091, IQR -0.041–0.251), indicating that alleles favored under stronger CAS selection were partly favored under weaker selection, but not the reverse. In both CAS candidate sets, β values under Low and High CAS differed significantly from those in Controls (Holm-adjusted *p* < 0.001), although the shift of Low CAS candidates under High CAS was minimal in magnitude.

**Figure 2.**
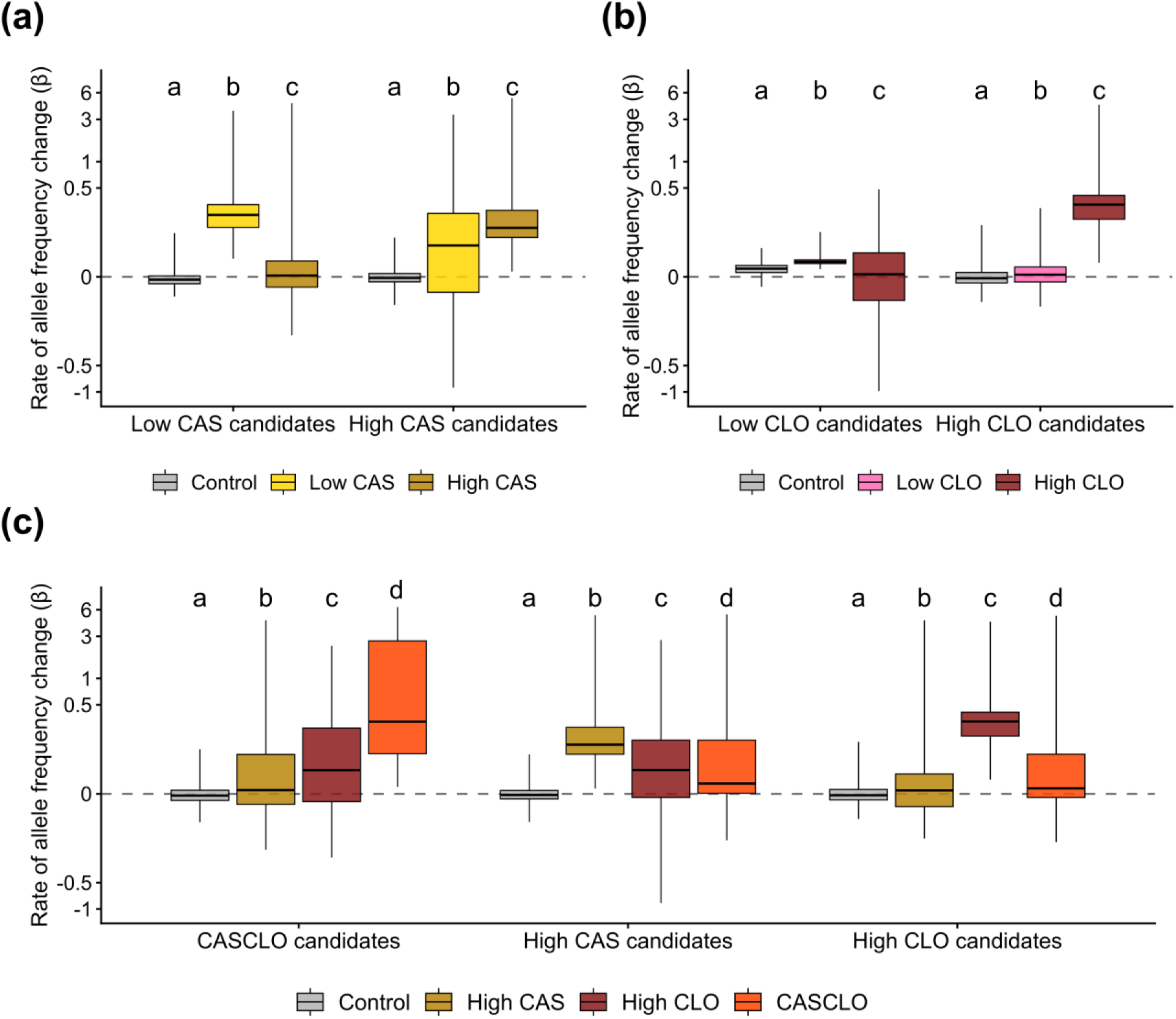
Rates of allele frequency change did not scale predictably with drug concentration or combination. Boxplots show the distribution of GLM effect sizes (β), representing rates of allele frequency change on the logit scale, with medians indicated by center lines. β was estimated using quasibinomial GLMs for each SNP. Panels show candidate SNPs from comparisons of CAS concentrations (a), CLO concentrations (b), and single-versus combined-drug treatments (c). SNPs identified as candidates in one treatment (top 5% by FDR-adjusted *p*-value) were evaluated across all related treatments, including Controls. For each SNP, β was polarized to the allele that increased in the treatment where the SNP was identified; positive values therefore indicate increases of the same allele in other treatments. Letters indicate significant differences among treatments (Friedman and pairwise Wilcoxon tests, Holm-adjusted *p* < 0.05). The y-axis was transformed using the inverse hyperbolic sine to display the full range of values while retaining resolution near zero.

CLO candidates showed a similar response. High CLO candidates shifted strongly under High CLO (β̃ = 0.320, IQR 0.213–0.411) but minimally under Low CLO (β̃ = 0.006, IQR −0.014–0.026; **Fig. 2b**). Low CLO candidates shifted only modestly under Low CLO (β̃ = 0.040, IQR 0.035–0.045) and even less under High CLO (β̃ = 0.007, IQR −0.065–0.067), and their response under Low CLO was only slightly greater than in Controls (β̃ = 0.021, IQR 0.011–0.030). This finding was consistent with the weaker signal in Low CLO populations observed in the PCA and SFS (**Figs. S2b and S3**).

We then tested whether combined drug exposure produced the additive response expected if each drug’s effects simply accumulated, in which case alleles favored under each single drug should also increase under CASCLO. Contrary to this expectation, single-drug candidates generally responded weakly under combined exposure. High CAS candidates shifted little under CASCLO (β̃ = 0.027, IQR 0.001–0.189), which was even less than under High CLO (β̃ = 0.066, IQR −0.009–0.189; **Fig. 2c**). High CLO candidates likewise shifted little under CASCLO (β̃ = 0.014, IQR −0.010–0.122) or High CAS (β̃ = 0.009, IQR −0.034–0.054). CASCLO candidates instead responded most strongly under combined selection (β̃ = 0.318, IQR 0.123–2.668), with much weaker shifts under High CAS (β̃ = 0.010, IQR −0.028–0.121) and High CLO (β̃ = 0.065, IQR −0.020–0.267). The median CASCLO response at these loci was roughly four times the sum of their responses under each drug alone (0.318 vs. 0.075), consistent with a non-additive response.

### Related antifungal selection regimes favor distinct sets of candidate loci

The β̃ comparisons showed that alleles favored in one treatment changed little in related environments. We therefore asked whether these environments nonetheless favored overlapping genomic targets. Because shared statistical significance alone could include loci with different allele frequency trajectories, we defined shared responses based on both statistical support and similarity in the rate of allele frequency change between treatments (see Methods). This allowed us to distinguish consistent responses from loci that were significant across treatments but followed different trajectories.

Because both drugs act on well-defined cellular targets, stronger stress might have been expected to draw on the same variants favored under weaker stress. Instead, candidate loci were largely treatment-specific. Under CAS selection, responding loci were distributed broadly across the genome in both Low and High CAS populations, with limited overlap between treatments (**Fig. 3a**). Among CAS candidates 2,010 SNPs were specific to Low CAS, 1,267 were specific to High CAS, and only 111 were shared between the two environments (**Fig. 3a**). The CLO comparison showed an even stronger separation among treatments, although interpretation was constrained by the limited genomic response under Low CLO. High CLO populations exhibited widespread genomic responses, whereas few loci showed responses specific to Low CLO. Among CLO candidates, 2,785 SNPs were specific to High CLO and only 66 were specific to Low CLO, with no loci showing shared responses between treatments (**Fig. 3b**).

**Figure 3.**
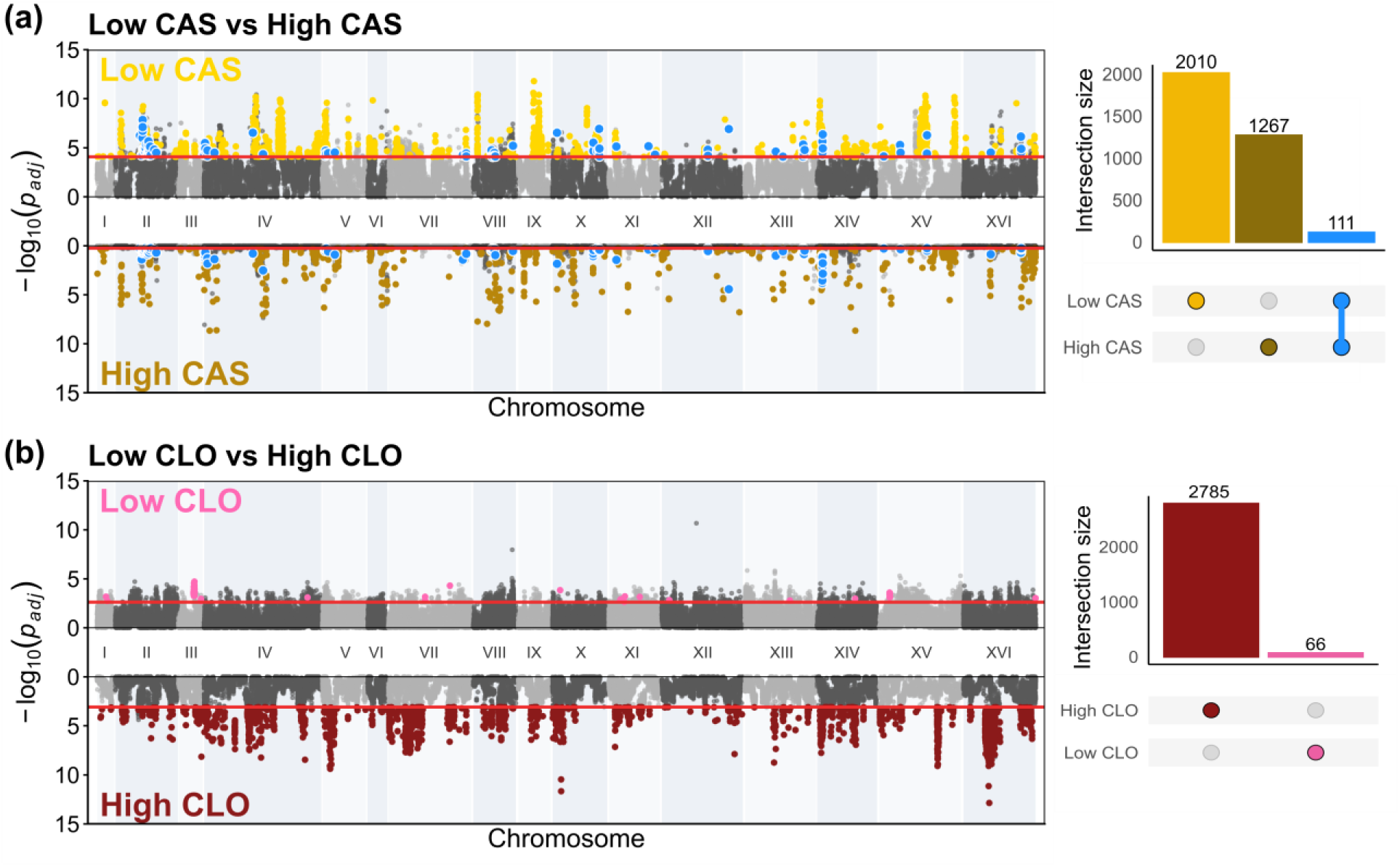
Loci responding to selection are largely distinct between populations evolved under different drug concentrations. Each panel shows a mirrored Manhattan plot paired with an UpSet plot for populations evolved under different concentrations of CAS (a) and CLO (b). In the Manhattan plots, each point represents a single SNP plotted by genomic position and −log_10_ FDR-adjusted *p*-value (y-axis), derived from quasibinomial GLMs fit to allele frequency trajectories across four timepoints (T_0_, T_1_, T_7_, T_14_). Points are shaded in alternating light and dark gray by chromosome. Red horizontal lines mark the significance threshold for each treatment, drawn at the 5^th^ percentile of FDR-adjusted *p*-values within that treatment. Candidate SNPs above this threshold are colored by their classification from pairwise slope contrasts of cross-treatment GLMs: gold (Low CAS-specific), dark gold (High CAS-specific), blue (shared between Low and High CAS) in (a); pink (Low CLO-specific) and dark red (High CLO-specific) in (b). UpSet plots summarize overlap among treatments, where vertical bars indicate the number of candidate SNPs in each intersection and the dots below identify the treatment combinations represented by each bar. No shared SNPs are shown in (b) because none met the criteria for a shared response between CLO concentrations.

Combined- and single-drug selection (CASCLO vs. High CAS and High CLO) also favored largely distinct loci. CASCLO populations showed strong candidate regions that did not overlap with the strongest single-drug responses, while broader regions responding under single-drug selection were reduced or absent under combined exposure (**Fig. 4**). Across candidate SNPs, 992 were specific to CLO, 322 were specific to CASCLO, and 229 were specific to CAS. The greatest overlap occurred between the two single-drug environments, where 147 SNPs responded to both CAS and CLO but not CASCLO. Under an additive expectation, these loci should have been among the most likely to respond to CASCLO exposure, suggesting that combining the drugs altered which loci were favored rather than accumulating the responses to each drug. In contrast, overlap between CASCLO and either single-drug environment was limited (61 with CAS, 60 with CLO), and only five SNPs showed shared responses across all three environments.

**Figure 4.**
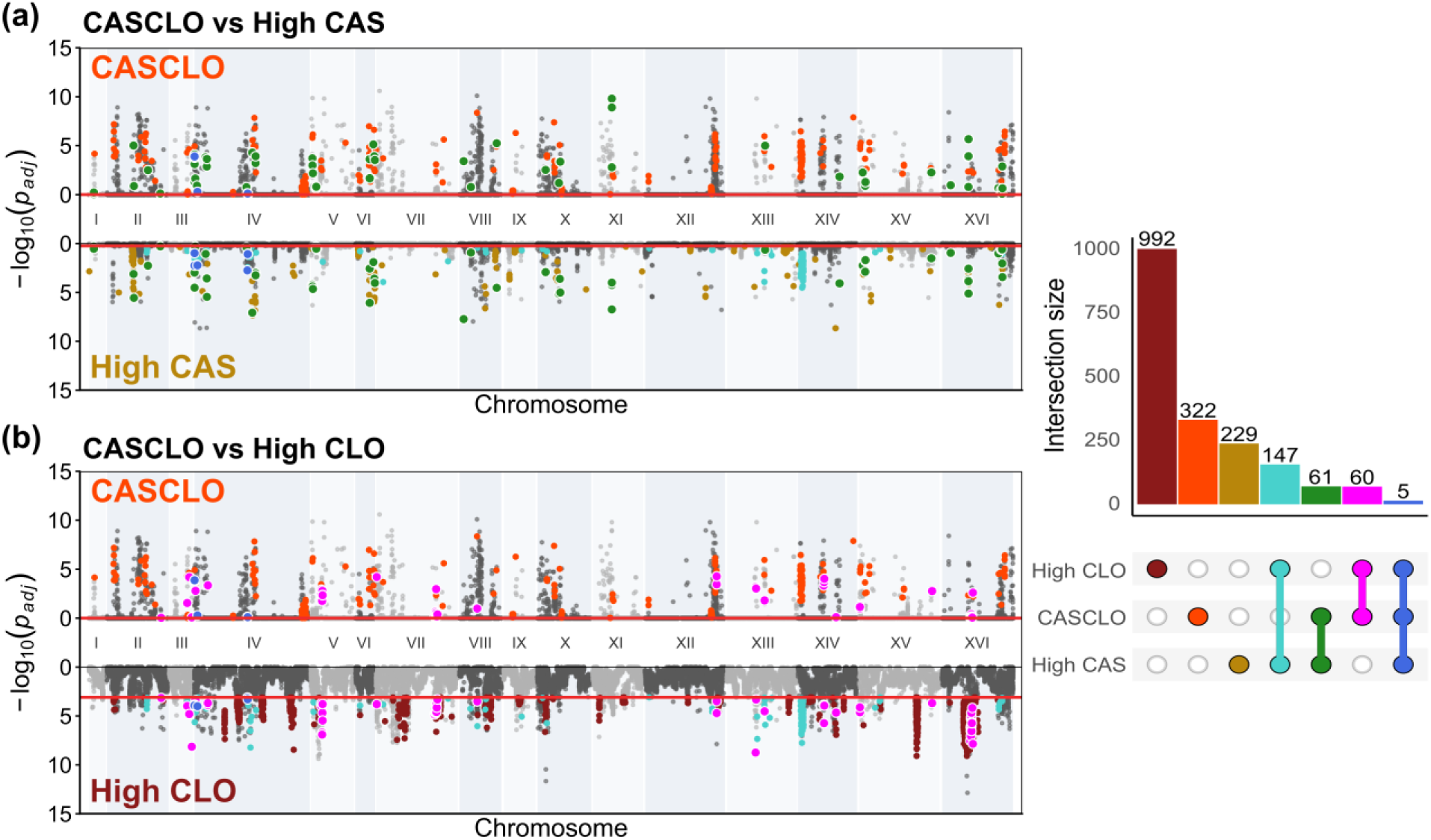
Combined drug treatment selects on largely distinct genomic targets from either drug alone. Each panel shows a mirrored Manhattan plot for populations evolved under combined drug treatment (CASCLO) compared with a single-drug treatment, High CAS (a) and High CLO (b), paired with an UpSet plot summarizing overlap across all three treatments. In the Manhattan plots, each point represents a single SNP plotted by genomic position (x-axis) and −log10 FDR-adjusted *p*-value (y-axis), derived from quasibinomial GLMs fit to allele frequency trajectories across four timepoints (T_0_, T_1_, T_7_, T_14_). Points are shaded in alternating light and dark gray by chromosome. Red horizontal lines mark the significance threshold for each treatment, drawn at the 5^th^ percentile of FDR-adjusted *p*-values within that treatment. Candidate SNPs above this threshold are colored by their classification from pairwise slope contrasts of cross-treatment GLMs: dark gold (High CAS-specific), dark red (High CLO-specific), orange (CASCLO-specific), dark green (High CAS & CASCLO shared), magenta (High CLO & CASCLO shared), turquoise (High CAS & High CLO shared), and royal blue (shared across all three treatments). The UpSet plot summarizes overlap among treatments, where vertical bars indicate the number of candidate SNPs in each intersection and the dots below identify the treatment combinations represented by each bar.

### Adaptation was largely polygenic, although known drug targets responded under specific conditions

To determine whether stronger or combined antifungal selection was more likely to engage previously characterized resistance targets, we annotated all candidate SNPs, including the early-fixed sites, using SnpEff (Cingolani et al. 2012) and summarized variants at the gene level. Because Low CLO populations showed limited genomic differentiation from Controls, gene-level comparisons focused on CAS populations (Low CAS vs. High CAS) and on combined-versus single-drug populations evolved under the strongest selection regimes (CASCLO vs. High CAS and High CLO). Candidate variants were distributed broadly among synonymous, regulatory, and missense changes, with high-impact variants rare (**Tables S2 and S3**).

We then cross-referenced candidate genes against the *S. cerevisiae* subset of the FungAMR database (Bédard et al. 2025), which compiles experimentally supported antifungal resistance genes. Overall, known resistance genes made up a small fraction of candidates and were not enriched in any gene set. Among CAS candidate genes, 23 of 1,008 candidate genes (2.3%) overlapped with FungAMR genes, with no significant enrichment in the Low CAS-specific, High CAS-specific, or shared CAS gene sets (all *p* > 0.05; Fisher’s exact test).

Similarly, only 15 of 691 candidate genes (2.2%) from the single-versus combined-drug comparison overlapped with FungAMR genes, with no significant enrichment detected in any gene set (all *p* > 0.05). Most FungAMR-associated variants were synonymous or non-coding (**Tables S4 and S5**). Stronger and combined selection were therefore not generally more likely to engage known resistance determinants.

Known targets nonetheless responded strongly in specific drug environments (**Fig. 5a**). The most relevant CAS-associated signal occurred at *FKS1*, which encodes the (1,3)-β-D-glucan synthase subunit targeted by CAS and is the primary determinant of echinocandin resistance (Katiyar and Edlind 2009). *FKS1* contained a dense cluster of candidate variants (187 synonymous, 15 missense) distributed across the length of the gene (∼5 kb) on chromosome XII. Three synonymous variants fell within the characterized echinocandin resistance hotspots, two in HS1 (residues 641–649) and one in HS2 (residues 1345–1365) (Perlin 2007; Johnson et al. 2011), whereas all missense candidates occurred outside these regions (**Fig. S4)**. At *FKS1* missense sites, the alternative allele segregated at intermediate frequency in the ancestor but was lost by T_1_ and remained absent through T_14_ in Low CAS, High CAS, and CASCLO populations. In contrast, this allele was maintained near ancestral frequency in Controls and declined more gradually under High CLO, reaching ∼0.1 by T_14_ (**Fig. 5b**). Selection at *FKS1* was thus specific to CAS exposure but independent of CAS concentration. Other CAS-associated FungAMR hits contained a mixture of synonymous and missense variants and likely reflected broader drug-stress responses or linkage with selected regions (**Tables S4 and S5**).

**Figure 5.**
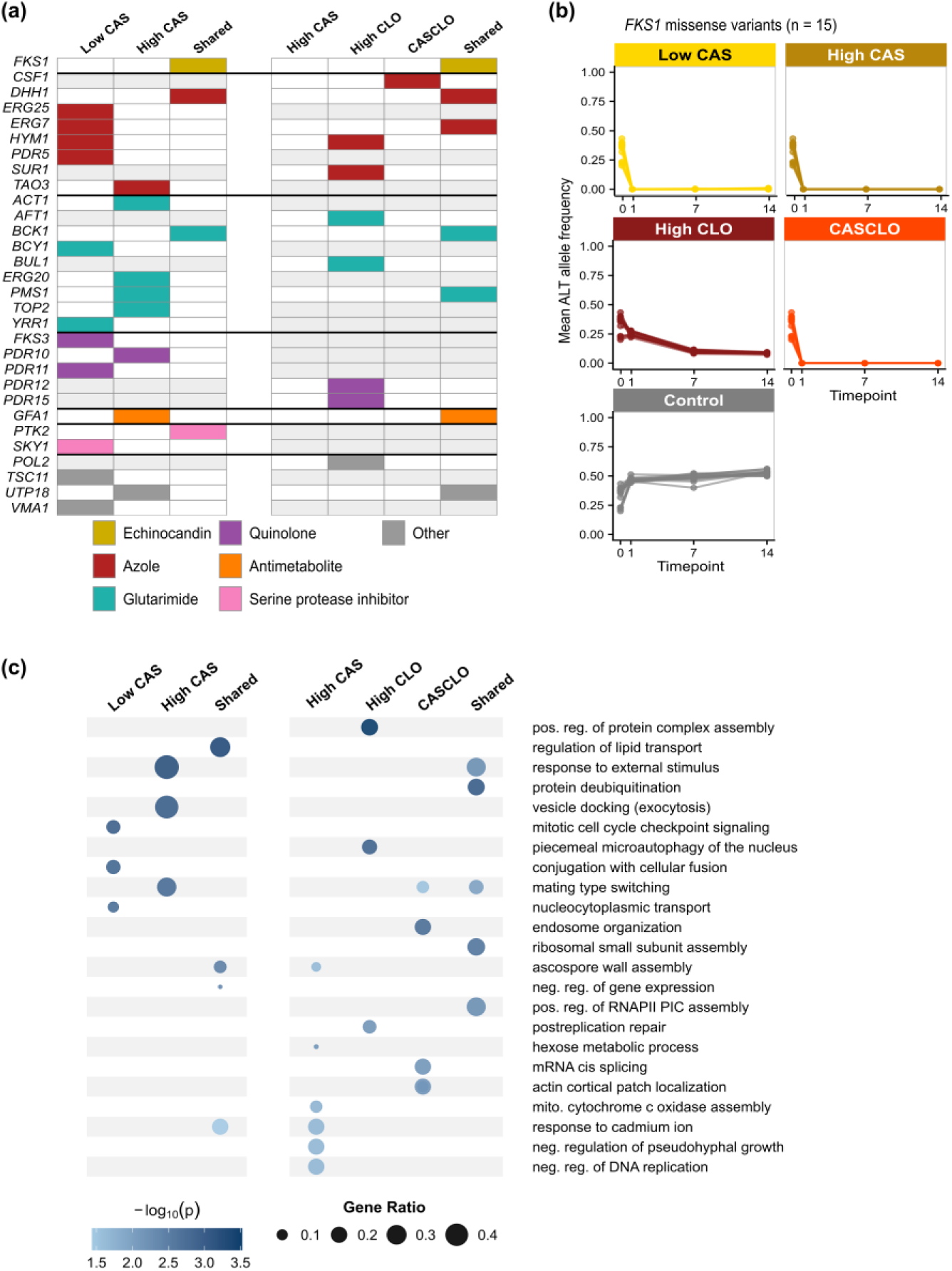
Resistance-associated genes and enriched functions differed across related drug treatments. (a) Candidate genes with known antifungal resistance associations in the FungAMR database (Bédard et al. 2025). Columns show gene categories from the CAS intensity comparison (Low CAS, High CAS, Shared), and the single-versus combined-drug comparison (High CAS, High CLO, CASCLO, Shared). Colored tiles indicate FungAMR hits by drug class. (b) Mean ALT allele frequency trajectories of *FKS1* missense candidate sites across 12 replicate populations; T_0_ is the ancestral value. (c) The three most significantly enriched GO biological process terms per gene set (topGO weight01 Fisher’s exact test, *p* < 0.05) after semantic reduction with rrvgo (Sayols 2023). Point size indicates gene ratio and color indicates −log₁₀(*p*). Full GO results are in Tables S6 and S7.

Fewer FungAMR genes responded under CLO or CASCLO selection. *HYM1* and *SUR1* were specific to CLO populations, *CSF1* was specific to CASCLO, and *ERG7* and *DHH1* were shared with single-drug environments (**Fig. 5a**). Most variants in these genes were synonymous, although *HYM1* and *SUR1* contained a small number of missense variants specific to CLO (*n* = 2 for *HYM1*, *n* = 1 for *SUR1*; **Table S5**) that showed the strongest allele frequency shifts in CLO populations, with similar directional changes under CASCLO (**Fig. S5**). Both genes have previously been implicated in azole adaptation (Song et al. 2008; François et al. 2009; Taylor et al. 2022); however, major azole resistance genes such as *PDR1*, *PDR3*, and *ERG11* (Mizoguchi et al. 2002; Diba et al. 2020) were not identified as candidates in any treatment.

Azole adaptation can proceed through copy number changes, most often amplification of the chromosome or region containing the drug target gene *ERG11* (Coste et al. 2007; Todd et al. 2023; Vande Zande et al. 2023). Because allele frequency analysis cannot detect such changes, we examined normalized sequencing coverage across the genome in all populations. Coverage of chromosome VIII, which carries *ERG11* in *S. cerevisiae*, increased under High CLO selection, rising from ancestral frequencies at T_1_ to ∼1.5× coverage by T_14_ in 9 of 12 replicate population_s_, while the remaining three stayed at ancestral frequencies (**Fig. S6a**). Coverage was elevated uniformly along the chromosome, providing putative support for whole-chromosome rather than segmental duplication (**Fig. S6b**). No comparable increase occurred in Control, Low CLO, or CAS populations. CASCLO populations showed replicate-specific variation in chromosome VIII coverage but no consistent directional increase, with coverage returning near ancestral levels by T_14_. However, sequencing coverage is an indirect measure of copy number, particularly in pooled samples where it reflects an average across many cells. We therefore interpret this pattern as a likely gain of chromosome VIII in at least a subset of cells within High CLO populations, consistent with a known route to azole resistance, but direct confirmation would require karyotyping.

### Functional enrichment was largely treatment-specific

To identify broader functional patterns, we performed gene ontology (GO) enrichment analysis on treatment-specific and shared candidate gene sets using topGO (Alexa and Rahnenführer 2025), with semantic reduction in rrvgo (Sayols 2023). We identified 11, 6, and 10 parent terms for Low CAS, High CAS, and shared CAS candidates, respectively. In the single-versus combined-drug comparison, we identified 7, 8, 6, and 9 parent terms for CAS-specific, CLO-specific, CASCLO-specific, and shared candidates, respectively (**Tables S6 and S7**). As with the candidate loci themselves, enriched functions were largely specific to individual gene sets, with only one term shared among gene sets within each comparison.

The most significantly enriched terms spanned diverse cellular processes (**Fig. 5c**), including mitotic cell cycle checkpoint signaling in Low CAS candidates (12 of 81 annotated genes; *p* = 0.001), response to external stimulus in High CAS candidates (5 of 11; *p* = 0.0007), and regulation of lipid transport in shared CAS genes (4 of 13; *p* = 0.0005). In the single-versus combined-drug comparison, the top terms were hexose metabolic process for CAS-specific candidates (5 of 91; *p* = 0.007), positive regulation of protein-containing complex assembly for CLO-specific candidates (7 of 33; *p* = 0.0003), and endosome organization for CASCLO-specific candidates (4 of 20; *p* = 0.002).

Given the known targets of CAS and CLO, we also looked for enriched terms related to cell wall and sterol processes. Several were enriched, though most did not rank among the top terms, and they occurred mainly in gene sets shared across treatments. Fungal-type cell wall organization was enriched in genes shared between Low and High CAS (15 of 172; *p* = 0.035) and in CAS-specific candidates from the single-versus combined-drug comparison (11 of 172; *p* = 0.026). Genes under this term included *FKS3*, a paralog of *FKS1* (Ishihara et al. 2007), and *GAS2* and *GAS3*, which encode glucan remodeling enzymes (Rolli et al. 2011). Genes shared between CASCLO and the single-drug treatments were enriched for (1,3)-β-D-glucan biosynthetic process (3 of 12; *p* = 0.007), the process CAS directly inhibits, and included *FKS1* and its negative regulator *LRG1* (Svarovsky and Palecek 2005). This shared gene set was also enriched for sterol import (2 of 10; *p* = 0.046), consistent with the membrane target of CLO.

### Phenotypic consequences of antifungal selection reveal environment-specific benefits and limited costs

To assess the phenotypic consequences of adaptation under different antifungal environments, we compared growth performance of evolved populations and the ancestor across control and drug-containing conditions. Growth was quantified using doubling time (DT) and carrying capacity (*K*) as proxies for fitness, capturing population growth rate and maximum density, respectively. We then asked whether stronger or combined selection improved performance under drug stress, and whether these gains came with costs under non-selective conditions.

Populations evolved under stronger drug selection showed improved growth (lower DTs) when challenged with the corresponding high-drug environment. High CAS populations grew faster than Low CAS populations when assayed in High CAS media (*p* < 0.05; **Fig. 6a**), and High CLO populations similarly outperformed Low CLO populations in High CLO media (*p* < 0.05; **Fig. 6b**). Relative to the ancestor, evolved populations also showed improved growth under high-drug conditions, with the ancestor failing to grow in CAS media and exhibiting substantially slower growth in High CLO media (**Fig. S7a, b**). Carrying capacity showed more treatment-specific patterns. High CLO populations reached higher *K* than Low CLO populations and the ancestor under High CLO conditions (*p* < 0.001; **Figs. 6b and S7d**), whereas CAS-evolved populations did not differ significantly in *K* (**Fig. 6a**). Notably, these consistent phenotypic gains arose despite the largely distinct genomic responses described above.

**Figure 6.**
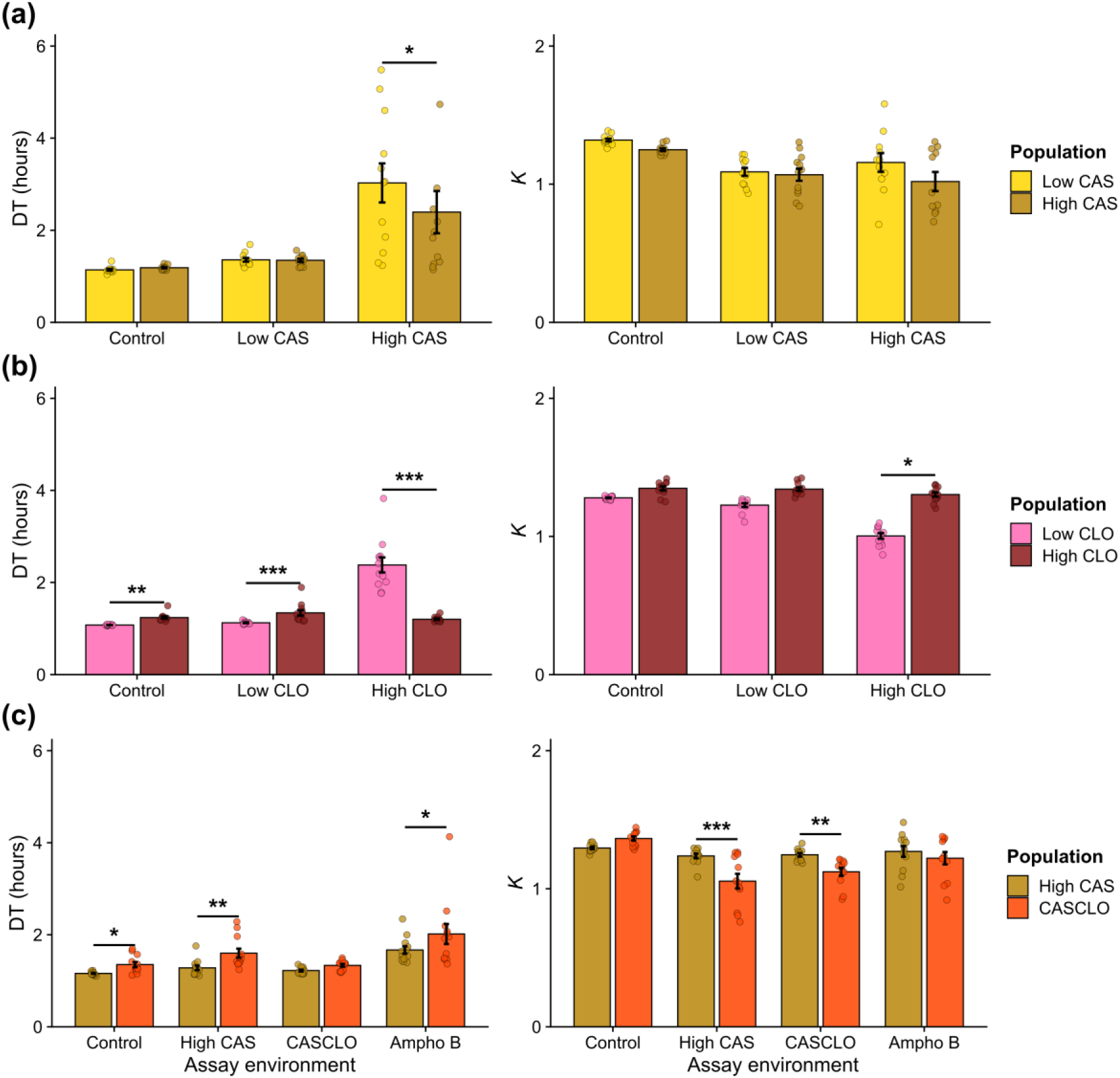
Growth of evolved populations was context dependent across drug environments. Doubling times (DT) and carrying capacities (*K*) were measured from 48 hr growth assays of evolved populations from the final timepoint (T_14_) across control and drug-containing media. Comparisons are shown for CAS selection (a), CLO selection (b), and CASCLO versus High CAS selection (c), with DT in the left column and *K* in the right column. Within each panel, the x-axis gives the assay environment and bars give the populations tested in that environment. DT and *K* values are shown as mean ± SE across replicate populations. Statistical significance was determined using linear mixed-effects models applied to log-transformed DT and *K* values, with replicate included as a random effect, and pairwise contrasts were Tukey-corrected. Asterisks indicate significant pairwise differences between populations within each assay environment (\**p* < 0.05; \*\**p* < 0.01; \*\*\**p* < 0.001).

Adaptation to higher drug concentrations did not produce substantial fitness costs under non-selective or milder conditions. Across control and low-drug conditions, DT and *K* were generally similar among evolved populations. High CLO populations were the only exception, showing slightly slower growth than Low CLO populations in control and Low CLO media (*p* < 0.01 and *p* < 0.001; **Fig. 6b**), but these differences were small in magnitude (ΔDT: ∼0.16–0.21 hr) and unlikely to represent major costs. Evolved populations also grew comparably to the ancestor under control conditions (**Fig. S7**), indicating that antifungal selection did not broadly reduce baseline fitness.

Combined-drug selection improved performance in the combined environment but did not provide a general advantage over single-drug selection. CASCLO populations grew in CASCLO media, where the ancestor failed to grow (**Fig. S7e**). However, CASCLO and High CAS populations had comparable DT in CASCLO media, and CASCLO populations grew more slowly and reached lower *K* than High CAS populations in control and High CAS media (**Fig.6c**).

Because adaptation to one antifungal can alter tolerance to others through shared resistance mechanisms (Lee et al. 2023), we also assayed growth in amphotericin B (AmphoB), a drug absent from the selection regime that targets membrane sterols through a distinct mechanism from CAS and CLO. Both CASCLO and High CAS populations grew in AmphoB media at rates comparable to those in their own selective environments (**Fig. 6c**). However, CASCLO populations grew more slowly than High CAS populations under AmphoB (DT: 1.92 vs. 1.65 hr; *p* < 0.05), suggesting that combined drug selection did not promote broader antifungal tolerance than adaptation to High CAS alone.

## Discussion

Experimental evolution has shown that populations often respond repeatably to similar selective conditions, but this predictability may decline as environments become more variable or complex (Bailey et al. 2015; Blount et al. 2018). We tested whether adaptive responses remain predictable as antifungal selection changes in intensity and complexity by evolving outcrossed *S. cerevisiae* populations under two drugs with distinct cellular targets, at low and high concentrations and in combination. Our results show that responses were shaped primarily by the specific selective environment rather than by drug concentration or number alone. Increasing drug concentration did not simply amplify the genomic response observed under weaker selection. Instead, it altered which variants responded, most clearly under CAS. Likewise, combined-drug selection produced a genomic response that was largely distinct from that under either drug alone, rather than the additive response expected if each drug’s effects accumulated. Known resistance mechanisms contributed in specific environments, including selection at *FKS1* under all CAS-containing treatments and a putative gain of the *ERG11*-carrying chromosome VIII under High CLO, but most candidate loci were environment-specific and not enriched for previously characterized resistance genes. Despite these divergent genomic responses, populations generally improved growth under their respective selective conditions, with limited evidence of fitness costs in the traits measured. Phenotypic outcomes were therefore more predictable than the genetic changes underlying them. This distinction is important for predicting evolution under changing antifungal environments, where repeatable phenotypic responses may conceal substantial genetic contingency. Overall, these findings highlight how changes in the intensity and combination of selective pressures can redirect adaptation toward different genetic solutions.

### Stronger selection largely redirects rather than amplifies a shared genomic response

Models of polygenic adaptation predict that stronger selection should produce larger allele frequency changes as populations move toward a more distant phenotypic optimum (Jain and Stephan 2017; Stetter et al. 2018; Christodoulaki et al. 2019; Thornton 2019). Because these models typically assume additive allele contributions, increased selection intensity should primarily amplify responses at variants already favored under weaker selection. However, the effects of genetic variants can depend on the environment in which they were selected, such that changing drug concentrations may alter which variants provide the greatest benefit (Gulisija 2026). Our results support this latter possibility. Although stronger antifungal selection produced more extreme allele frequency changes at some sites, it did not consistently amplify responses at the same loci. Instead, the variants that responded largely differed between drug concentrations, suggesting that higher drug concentrations shifted the targets of selection.

If stronger antifungal selection primarily amplified existing responses, alleles favored under low drug concentrations should have shifted further under high concentrations. Instead, the relationship between CAS treatments was asymmetric. Alleles favored under High CAS were partly favored under Low CAS, shifting at roughly half their High CAS rate, whereas alleles favored under Low CAS changed little under High CAS (**Fig. 2a**). Some variants favored under strong CAS selection may therefore also have provided benefits at lower concentrations, whereas variants favored under weaker selection may have offered little additional benefit as stress intensified. Together with the limited overlap among candidate loci (**Fig. 3a**), these results indicate that Low and High CAS populations followed largely distinct genomic trajectories, even though both showed strong selection against the alternative *FKS1* allele (**Fig. 5b**).

Drug concentration has similarly been shown to alter the genetic basis of resistance in other microorganisms. In *Candida albicans*, lineages evolved to different fluconazole concentrations acquired different mutational spectra (Todd et al. 2023), and in *Escherichia coli*, different antibiotic exposure levels favored distinct genetic solutions (Harmand et al. 2017).

Those studies examined new mutations arising in clonal populations, whereas our results reveal a similar pattern during adaptation from standing genetic variation. This variation can provide a major source of rapid adaptation in genetically diverse populations, including clinical fungal pathogens (Burrack et al. 2022). One possibility is that increasing drug stress changes the relative benefits of existing variants, favoring those that improve survival under greater physiological constraint while balancing the costs of resistance (Melnyk et al. 2015). Different combinations of alleles may also provide similar levels of tolerance, allowing populations to adapt through different genetic routes (Barghi et al. 2020). Such genetic redundancy could help explain why High CAS populations remained broadly polygenic rather than converging on a small number of strongly-responding loci.

Stronger antifungal selection also increased the number of sites that reached extreme allele frequencies early in the experiment, consistent with stronger directional selection driving more rapid allele frequency changes (Christodoulaki et al. 2019). However, these sites also differed substantially among treatments, with most specific to one treatment. Stronger and combined selection therefore produced larger allele frequency changes, but at a largely different set of sites, reinforcing that the selective environment shaped which loci responded and how far they moved.

CLO adaptation showed a similar pattern, although the evidence was less clear at low concentrations. Low and High CLO had no overlapping candidate loci (**Fig. 3b**), and a putative gain of chromosome VIII, which carries *ERG11*, was observed only under High CLO (**Fig. S6**). However, we interpret these differences cautiously because Low CLO populations showed minimal genomic differentiation from Controls (**Fig. S2b**), and candidate alleles shifted only slightly more under Low CLO than in Controls (β̃ = 0.040 vs. 0.021; **Fig. 2b**). These results raise the possibility that some observed changes reflected laboratory adaptation rather than a response to Low CLO itself. The ancestor was also more tolerant of CLO than CAS, making Low CLO a comparatively weak selective challenge over the experimental timeframe, despite preliminary work indicating moderate effects of the Low CLO dosage (**Fig. S1**). The limited genomic response under Low CLO may therefore reflect a stress level too low to produce detectable allele frequency shifts, rather than a fundamentally distinct adaptive trajectory. Nevertheless, Low CLO populations grew better than the ancestor under High CLO conditions (**Fig. S7c**), providing some evidence that phenotypic adaptation occurred even though its genomic signature was difficult to distinguish from background change.

### Combined drug exposure favors a distinct, non-additive genomic response

If the effects of CAS and CLO were simply additive, we would expect genomic responses in CASCLO populations to largely overlap with the responses observed under each drug alone, but perhaps at greater magnitude as their contributions accumulated. Instead, candidate loci favored under single-drug exposure generally shifted little under CASCLO, and variants favored under High CAS shifted more under CLO alone than under the combined treatment (**Fig. 2c**). In contrast, variants favored under CASCLO showed a median response roughly four times the sum of their responses under the individual drugs. Candidate loci also overlapped more between the two single-drug treatments than between either single drug and CASCLO, with few loci shared across all three environments (**Fig. 4**). Although CASCLO combined the drugs at lower concentrations, its genomic response differed from those under High CAS and High CLO, which also strongly inhibited ancestral growth (**Fig. S1**). This comparison does not directly control for differences in selection strength, but the results suggest that combined exposure favored different genetic changes rather than simply intensifying responses to either drug alone.

One explanation is that exposure to both drugs altered the relative benefits of individual alleles. Variants that were advantageous under a single drug may have offered less benefit under combined exposure, while other alleles became more important under the simultaneous stress on cell wall and membrane function. We cannot determine the exact mechanisms underlying this shift from our data, but non-additive responses to multiple stressors have also been reported in *Daphnia* (Cuenca-Cambronero et al. 2021), *Drosophila* (Burny et al. 2022), marine copepods (Brennan et al. 2022), and *S. cerevisiae* (Rêgo et al. 2024). Similar patterns have emerged in bacterial evolution experiments, where drug combinations redirected resistance evolution by changing which variants provided the greatest benefit under combined selection (Hickman et al. 2017; Jahn et al. 2021). Notably, this genomic divergence contrasts with the pharmacological effects of echinocandin-azole combinations, which often show synergistic or additive inhibition of fungal growth (Petraitis et al. 2003; Khalifa et al. 2021; Toepfer et al. 2024). Thus, predictable drug interactions at the growth level do not necessarily translate into predictable evolutionary responses, limiting the extent to which single-drug experiments can inform predictions of adaptation under multidrug regimens.

Despite these differences, CASCLO populations retained some features of the response to CAS. Most variants that fixed rapidly under CASCLO were also fixed under High CAS, including variants at *FKS1*. This overlap suggests that changes at the primary echinocandin target remained important even in the presence of CLO. Genes shared between CASCLO and the single-drug treatments were also modestly enriched for processes tied to both drug targets, including (1,3)-β-D-glucan biosynthesis and sterol import. This overlap may reflect the close relationship between the cell wall and plasma membrane, where disruption of one can affect the physiological consequences of perturbing the other (Levin 2005; Fabri et al. 2020). In contrast, CASCLO-specific genes were enriched for other processes, including endosome organization and mRNA cis splicing (**Fig. 5c**). Although the role of these processes remain unclear, their enrichment indicates that adaptation to combined drug exposure involved cellular processes beyond those associated with either drug alone.

### Antifungal adaptation combined target-associated changes with broader genomic responses

Although antifungal resistance is often defined by a small number of repeatedly implicated genes, these loci represent only a subset of the mechanisms that can influence survival under drug exposure (Cowen et al. 2015; Lee et al. 2023). In our populations, candidate variants were distributed throughout the genome, with relatively little overlap with established resistance genes. The clearest exceptions were selection at *FKS1* under CAS exposure and increased chromosome VIII coverage under High CLO. Beyond these responses, candidate loci and enriched functional categories differed across treatments and reflected broader cellular processes that varied with drug identity, concentration, and combination (**Fig. 5c**).

A similar pattern was reported in *S. cerevisiae* evolved under moderate and high ethanol stress, where the two intensities engaged partially distinct biological pathways (Phillips et al. 2026). Our results extend this observation to stressors with more specific cellular targets, and found that drug adaptation extended beyond the genes directly associated with those targets. *FKS1*, which encodes the primary target of echinocandins (Katiyar and Edlind 2009), showed the most consistent response across conditions, with rapid allele frequency changes under Low CAS, High CAS, and CASCLO (**Fig. 5b**). These changes arose from variants already present in the ancestral population rather than newly arising resistance mutations. Standing variation at a major drug target can therefore contribute to adaptation across distinct selective environments, even when the broader genomic responses differ.

The variants in *FKS1* also differed from the canonical echinocandin resistance substitutions typically associated with this gene. Most known echinocandin resistance mutations occur within two hotspot regions, HS1 and HS2 (Perlin 2007; Johnson et al. 2011). In our CAS populations, candidate variants were distributed more broadly across the locus (**Fig. S4**). This raises the possibility that CAS selection acted on a haplotype or linked genetic background rather than favoring individual substitutions independently. These variants could influence CAS susceptibility through linkage with functional variants or through effects on gene regulation or protein interactions, but functional characterization will be needed to determine whether these variants contribute directly to the observed response.

Outside *FKS1*, few candidate loci overlapped with established antifungal resistance genes. We did find some evidence for azole response genes under High CLO, including *SUR1* and *HYM1* (**Fig. S5**) (Song et al. 2008; François et al. 2009; Taylor et al. 2022), as well as a putative gain of chromosome VIII containing *ERG11*. Increased *ERG11* dosage has been linked to azole resistance in fungi, where additional copies of the drug target raise enzyme levels enough to offset partial inhibition while preserving the primary function of the ergosterol pathway (Coste et al. 2007; Todd et al. 2023; Vande Zande et al. 2023). In 9 of 12 High CLO populations, coverage of chromosome VIII was elevated uniformly rather than within a discrete segment (**Fig. S6**). A similar chromosome VIII gain was observed in CLO-evolved *S. cerevisiae* populations that lacked point mutations in *ERG11* (Taylor et al. 2022), and aneuploidy is a common and reversible response to azole stress in pathogenic fungi (Todd and Selmecki 2023). Thus, the putative gain of chromosome VIII in most High CLO populations may have provided an additional route to CLO adaptation, potentially through increased *ERG11* dosage.

Notably, we did not observe a comparable increase in coverage under CASCLO despite the presence of CLO. Increased *ERG11* dosage may have been favored under High CLO exposure but was less advantageous in the combined drug environment. The balance between increased *ERG11* dosage and the costs of aneuploidy could contribute to this difference, although the underlying mechanism remains unresolved.

### Drug tolerance improved with limited trade-offs

Evolutionary adaptation to antimicrobial stress often involves fitness trade-offs, where genetic changes improve survival in the presence of drugs but reduce performance in drug-free environments or under alternative stresses (Chait et al. 2016; Schmidlin et al. 2024). These costs may arise when resistance mechanisms alter normal cellular processes or impose energetic demands that are beneficial only during continued drug exposure (Melnyk et al. 2015). We therefore expected populations evolved under High CAS, High CLO, and CASCLO to show some loss of fitness under less stressful conditions.

Contrary to this expectation, evolved populations generally improved growth under drug exposure without a corresponding decline in drug-free conditions. High CAS and High CLO populations showed the strongest improvements in their corresponding high-drug environments, outperforming populations evolved under lower drug concentrations (**Figs. 6a, b**). For CAS, these gains reflected faster growth rather than increased carrying capacity, whereas High CLO populations improved in both. This contrasts with patterns reported in other yeast evolution studies, where faster growth has been associated with lower final population yield (Sezmis et al. 2018). Here, increased growth under drug stress did not generally come at the expense of carrying capacity.

One explanation for this pattern is compensatory evolution, in which secondary changes outside the primary drug targets offset the costs associated with resistance (Maisnier-Patin and Andersson 2004). We identified changes at several loci beyond *FKS1* and *ERG11*, although their contributions to fitness remain unclear. Such compensation is plausible given the nature of the antifungals. In response to echinocandins, for example, fungi can increase chitin synthesis to reinforce cell walls weakened by drug exposure (Walker et al. 2010). This response could help explain how CAS adaptation was maintained without an obvious loss of growth. Alternatively, some resistance mechanisms may impose little detectable cost under benign conditions (Bedard et al. 2024). In *C. albicans*, *ERG11*-associated aneuploidies have been linked with increased azole resistance without a detectable fitness penalty in the absence of drug (Selmecki et al. 2009). Similarly, the increased chromosome VIII coverage observed in our High CLO populations was not accompanied by a substantial reduction in growth under the conditions tested. These explanations are not mutually exclusive, and the absence of detectable costs in our assays does not rule out trade-offs in other traits or environments.

High CAS and CASCLO populations also grew in AmphoB, despite never being exposed to this drug during evolution (**Fig. 6c**). Although CAS and AmphoB act on different primary targets (Lee et al. 2023), adaptation to CAS-associated cell-wall stress may have altered broader cellular responses that also influence susceptibility to AmphoB. AmphoB exposure itself alters the expression of genes involved in both ergosterol synthesis and cell-wall biogenesis in *S. cerevisiae* (Zhang et al. 2002). We did not detect significant responses at genes previously implicated in AmphoB susceptibility (Young et al. 2003; Balkan et al. 2019; Kong et al. 2020), so this may reflect changes in processes shared across antifungal responses rather than a direct response at known loci.

## Conclusion

Our results show that adaptation to antifungal stress becomes more genetically distinct, rather than simply more pronounced, as drug selection intensifies and increases in complexity. This distinction is particularly relevant for antimicrobial evolution, where treatment conditions change over time and resistance can emerge through diverse genetic routes. More broadly, genomic responses observed under one selective environment may provide limited insight into adaptation under stronger or more complex stresses, even when some phenotypic outcomes remain similar. Experimental evolution studies should therefore more carefully consider the design of selection regimes when extrapolating findings to natural and clinical settings.

## Materials and Methods

### Antifungal drugs and concentration determination

Caspofungin acetate (CAS) and clotrimazole (CLO) were obtained from GoldBio (St. Louis, MO, USA), prepared as stock solutions in DMSO, filter-sterilized, and stored at -20°C prior to use. Drug concentrations for the evolution experiment were selected based on dose-response assays conducted under conditions mirroring the experimental evolution regime (see Supplementary Methods). Two concentrations per drug were selected to represent distinct levels of selective pressure: Low CAS (0.035 µM), High CAS (0.07 µM), Low CLO (0.35 µM), and High CLO (3.00 µM), with low concentrations producing modest inhibition and high concentrations producing substantial but sublethal inhibition, allowing populations to reach sufficient density for transfer within 24–48 hr. For the CASCLO combination, the low-concentration pairing (0.035 µM CAS + 0.35 µM CLO) was selected because higher combined concentrations produced complete growth arrest, with no measurable increase in cell density over a 48-hr incubation period (**Fig. S1**).

### Experimental evolution

Experimental populations were founded from the genetically diverse *S. cerevisiae* 4S ancestral population, an outcrossed and highly recombined population derived from four SGRP founder strains including YPS128, Y12, DBVPG6765, and DBVPG6044 (Cubillos et al. 2009). The 4S population was generated through additional outcrossing and strain engineering to increase recombination, maintain extensive standing genetic variation, and promote adaptation to laboratory conditions (Burke et al. 2020; Linder et al. 2020; Phillips et al. 2021). Further details regarding the 4S population are provided in **Supplementary Methods**.

The 4S ancestral population was revived from 1 mL of a frozen stock on YPD agar at 30°C for 48 hr. Cells were then used to initiate an overnight liquid culture in 10 mL YPD broth. The overnight culture was used to found 12 biological replicate populations per treatment across six selective conditions (Low CAS, High CAS, Low CLO, High CLO, CASCLO, and DMSO control), resulting in 72 total populations. These conditions varied selection along two dimensions: intensity, manipulated through drug concentration (low vs. high), and complexity, manipulated through single-versus combined-drug exposure. Starting population sizes were standardized to an OD_600_ ∼ 0.1 in 1 mL of the appropriate selective medium and added to alternating wells of 24-well plates (Corning), with intervening wells containing sterile YPD broth to monitor for potential cross-contamination between wells; no contamination was detected over the course of the experiment. Plates were incubated for 24 hr at 30°C with shaking at 200 rpm.

Following incubation, cell density was quantified by a Tecan Spark microplate reader (Tecan Group Ltd., Männedorf, Switzerland), and transfer volumes were calculated to passage ∼10^6^ cells into 1 mL of fresh selective medium at each transfer, minimizing bottleneck effects and maintaining a large effective population size throughout the experiment.

Daily transfers were performed for 14 days, resulting in 13 total transfers for most populations. Consistent with dose-response assays, populations evolving under High CAS and CASCLO treatments exhibited limited growth during their initial 24 hr exposure to drug selection (**Fig. S1**), yielding insufficient cell densities for transfer. These populations were therefore incubated for an additional 24 hr prior to their first transfer, resulting in 12 total transfers over the evolution period. At multiple timepoints, including after the first transfer (T_1_), mid-experiment (T_7_), and at the end of the experiment (T_14_), 1 mL of each population was collected and preserved at -80°C in 15% glycerol for downstream phenotyping and sequencing. T_1_ therefore corresponded to 24 hr of drug exposure for most populations and 48 hr for High CAS and CASCLO populations, although limited growth during the additional 24 hr meant that few additional generations elapsed before T_1_ (**Table 1**).

The number of generations per transfer was estimated from OD_600_-derived cell density measurements as log_2_(*N*_final_/*N*_initial_), where *N*_initial_ and *N*_final_ represent estimated cell numbers at the start and end of each 24 hr incubation period, respectively. Cumulative generations ranged from 54 to 69 across treatments (**Table 1**), reflecting differences in growth rates of populations under different drug concentrations or combinations.

### Pooled-population genome sequencing

Genomic DNA was isolated from populations archived at T_1_, T_7_, and T_14_, yielding 216 samples (12 replicate populations × 6 selection regimes × 3 timepoints). To prepare each sample, 700 µl of glycerol stock was revived on YPD agar plates at 30°C for 48 hr. Yeast lawns were broadly sampled with a sterile wooden applicator and inoculated into 10 mL of YPD broth, which was grown overnight at 30°C with shaking at 200 rpm. Approximately 1 mL of the resulting culture was used for DNA isolation using the Qiagen Gentra Puregene Yeast/Bact. Kit (Qiagen, Hilden, Germany; Cat. No. 158567) following the manufacturer’s protocol. Genomic DNA was quantified with a Qubit fluorometer (Thermo Fisher Scientific, Waltham, MA, USA), and sequencing libraries were prepared using a modified Nextera protocol (Illumina, San Diego, CA, USA) as described in the **Supplementary Methods**. Briefly, DNA was diluted and tagmented before PCR amplification with dual-index barcodes. Amplified libraries were cleaned, pooled, size-selected, and purified prior to sequencing on an Illumina NextSeq 2000 platform with a 2 × 150 bp paired-end setup at the Center for Quantitative Life Sciences (CQLS; Oregon State University, Corvallis, OR, USA). Low coverage samples were re-pooled and re-sequenced to obtain higher and more consistent coverage across all populations.

### Sequence processing, variant calling, and quality control

FASTQ sequence data were processed using a variant calling workflow based on GATK to generate a combined VCF file containing all detected single-nucleotide polymorphisms (SNPs) across replicates and treatments, using Nextflow (Di Tommaso et al. 2017) to increase efficiency and reproducibility. Briefly, raw reads were aligned to the *S. cerevisiae* S288C reference genome (R64-1-1) using BWA MEM v0.7.18 (Vasimuddin et al. 2019), and variants were called using GATK v.4.5.0.0 (Van der Auwera and O’Connor 2020) following best practices (Poplin et al. 2017). A reference VCF file containing known SNPs from 1,011 natural *S. cerevisiae* isolates (Peter et al. 2018) was used by GATK BaseRecalibrator and ApplyBQSR to calibrate base quality scores. The resulting VCF file was filtered with GATK VariantFiltration using the following parameters: “QD < 5.0, MQ < 40.0, FS > 60.0, ReadPosRankSum < -8.0, MQRankSum < -12.5”. Allele depth (AD; number of reads supporting the alternate allele) and total coverage (DP) fields were extracted from the merged VCF for all high-quality variants, and allele frequencies were calculated as AD/DP at each site.

The dataset was then filtered to retain high-quality, biallelic polymorphic SNPs. Sites with a read depth below 10 in any experimental population were excluded. To focus on the evolution of standing genetic variation, we retained only variants that were polymorphic in the ancestral population, defined as allele frequencies between 0.02 and 0.98. This filter also removed very low-frequency calls likely to reflect sequencing or variant calling error. Additional low-confidence sites were removed based on the founder genotypes, following Phillips et al. (2021). Because the four founder strains are haploid and isogenic, sites appearing polymorphic within a founder likely reflect mapping or sequencing error and were excluded. Sites at which all four founders carried the same allele were also excluded, since no standing variation could have contributed to the 4S population at these sites. After filtering, the final dataset consisted of 64,764 high-quality polymorphic SNPs with a median genome-wide coverage of 86× (range: 27×–290×; **Table S1**).

### Assessment of variation prior to modeling

To evaluate the structure and quality of the dataset before statistical modeling, allele frequency patterns were examined using principal component analysis (PCA) and site frequency spectra (SFS). Sequencing coverage across chromosome VIII was also examined for putative evidence of amplification involving the *ERG11* region, a known mechanism of azole resistance (Coste et al. 2007). Details of both analyses are provided in **Supplementary Methods**.

The SFS revealed a subset of variants at allele frequencies of 0 or 1, indicating fixation or loss of the reference or alternate allele in the pooled data. Many of these variants had already reached fixation at T_1_ and remained fixed thereafter. We therefore defined a set of early-fixed sites as variants that reached a frequency of 0 or 1 by T_1_ and remained fixed at T_7_ and T_14_ in every evolved replicate within a treatment. This definition captures only variants that fixed rapidly and consistently, and excludes sites that fixed later in the experiment or in a subset of replicates. Because these sites show no remaining temporal variance, they could not be included in the regression-based modeling and were instead characterized separately as a complementary set of candidate loci. Early-fixed sites were filtered to exclude those also fixed in Control populations, which likely reflect laboratory adaptation unrelated to drug exposure, and those showing minimal change from the ancestor (|Δallele frequency| < 0.1). Sites were then classified as treatment-specific or shared based on which treatments they fixed in.

### Statistical modeling of allele frequency trajectories

SNP-level data from the ancestral population (T_0_) and three evolutionary timepoints (T_1_, T_7_, T_14_) were fit using quasibinomial generalized linear models (GLMs) with the ‘glm’ function in R. Early-fixed sites, defined as variants reaching a frequency of 0 or 1 by T_1_ and remaining fixed thereafter in every replicate within a treatment, were excluded from modeling because the absence of temporal variation precludes regression-based inference; these sites were analyzed separately. For all remaining SNPs, the quasibinomial family was used to account for overdispersion common in pooled sequencing data, arising from variability in read depth and stochastic sampling. This approach follows Wiberg et al. (2017), who showed that quasibinomial GLMs improve control of false positives and provide more accurate inference of consistent allele frequency change compared with other commonly used tests. Within these models, sequencing coverage at each SNP was incorporated as a weighting factor to account for differences in read depth.

Two complementary modeling approaches were applied. First, separate models were fit for each treatment to test whether allele frequencies changed over time within that environment (*allele frequency ∼ time*), with time treated as a continuous predictor. Second, cross-treatment models were constructed to formally compare allele frequency change among treatments (*allele frequency ∼ time × treatment*). Three cross-treatment models were implemented to reflect different experimental contrasts: a CAS intensity model (Low CAS, High CAS, and Control), a CLO intensity model (Low CLO, High CLO, and Control), and a single-versus combined-drug model (High CAS, High CLO, CASCLO, and Control). The single-versus combined-drug model used the high single-drug treatments so that differences would more closely reflect the combination of drugs rather than selection strength alone. In each model, the *time × treatment* interaction term enabled statistical comparison of rates of allele frequency change among treatments. The rate of change within each treatment was estimated using the ‘emtrends’ function from the “emmeans” package v2.0.0 (Lenth and Piaskowski 2025), and pairwise comparisons of these slopes were performed using the ‘pairs’ function.

Slope estimates (β coefficients, logit scale) and *p*-values were extracted for each SNP from single-treatment models, and pairwise slope contrasts and associated *p*-values were extracted from cross-treatment models. *P*-values were adjusted for multiple testing using the false discovery rate (FDR) correction (Benjamini and Hochberg 1995). Candidate SNPs were then identified by applying a significance threshold at the 5^th^ percentile of FDR-adjusted *p*-values within each treatment, retaining the top 5% of SNPs with the strongest evidence of allele frequency change over time. To compare how these loci responded across treatments, β values were polarized to the allele that increased in the treatment where each candidate was identified, so that positive values in other treatments indicate change in the same direction. Polarized β values were compared among treatments using Friedman rank-sum tests (β ∼ Treatment | SNP) with paired Wilcoxon contrasts (Holm-adjusted) and summarized as median coefficients (β̃).

### Classification of treatment-specific and shared responses

To compare which loci responded across treatments, SNPs were classified using both statistical support and similarity in the rate of allele frequency change. A SNP was treated as a candidate in a given treatment if its FDR-adjusted *p*-value fell within the top 5% for that treatment, as defined above. Slopes were then compared among treatments using the pairwise contrasts from the cross-treatment models, with a threshold of *p* < 0.05.

SNPs were classified as treatment-specific when they were candidates in one treatment only and their rate of allele frequency change differed significantly from that in every other treatment, including Controls. SNPs were classified as shared when they were candidates in two or more drug treatments, their rates of change did not differ significantly among those treatments, and each differed significantly from Controls. Requiring a significant difference from Controls helped ensure that responses in drug-evolved populations reflected drug exposure rather than laboratory adaptation. Candidate SNPs were visualized as Manhattan plots of -log_10_ FDR-adjusted *p*-values and colored by classification in R. UpSet plots summarizing the number of SNPs in each treatment-specific and shared group were generated using the R package “ComplexUpset” v1.3.3 (Krassowski 2020).

### Functional annotation, resistance gene enrichment, and gene ontology analysis

All candidate SNPs, including early-fixed sites, were annotated using SnpEff v5.4a (Cingolani et al. 2012). SNPs were classified by predicted functional effect, including synonymous, missense, nonsense, and intergenic variants, and the highest-impact annotation was retained for each SNP using the SnpEff impact hierarchy (High > Moderate > Low > Modifier). Candidate SNPs were then collapsed to the gene level by compiling all genes associated with at least one candidate variant within each treatment category. Because Low CLO populations yielded limited candidate loci, drug intensity comparisons were restricted to CAS treatments. For the CAS comparison, genes containing candidate variants only in Low CAS populations were classified as Low, those only in High CAS as High, and those with variants in both as Shared.

For the single-versus combined-drug comparison, genes were similarly classified as High CAS, High CLO, CASCLO, or Shared. Because gene-level groupings were based on the presence of candidate variants rather than on the SNP-level groupings described above, a gene containing variants from more than one category was assigned to the Shared group.

These gene lists were queried against FungAMR (Bédard et al. 2025), a curated database of fungal antimicrobial resistance-associated genes and mutations, using a custom R script.

Candidate genes were compared against FungAMR entries to identify overlap with genes previously linked to antifungal resistance phenotypes, clinical resistance, or experimentally characterized resistance-associated variants. Fisher’s exact tests were used to assess whether each candidate gene category was enriched for known resistance genes relative to the full background of candidate genes, with *p*-values adjusted for multiple comparisons using the FDR correction.

The same gene lists were used for gene ontology (GO) enrichment analysis to identify biological processes associated with treatment-specific candidate variants. Candidate gene lists were restricted to recognized yeast ORFs to ensure reliable GO term mapping, and the gene universe for enrichment testing consisted of all genes containing at least one SNP in the dataset and mapped to a valid yeast ORF in the org.Sc.sgd.db database (*n*=5,940 genes). GO enrichment of biological process terms was tested separately for each gene category using “topGO” v2.62.0 (Alexa and Rahnenführer 2025) and “org.Sc.sgd.db” v3.22.0 (Carlson 2025), with the weight01 algorithm and Fisher’s exact test (minimum node size of 10 genes; *p* < 0.05). Significant terms were reduced for redundancy using “rrvgo” v1.22.0 (Sayols 2023) with the Wang semantic similarity measure and a similarity threshold of 0.9, and the most significant term within each group was retained as its parent term. Overlap in enriched functions among gene sets was assessed using these aggregated parent terms.

### Analysis of growth rate differences

We assessed growth performance of populations from the final timepoint (T_14_) and the ancestral population in control medium and in the drug media relevant to each selection regime. High CAS and CASCLO populations were additionally assayed in 2 µM amphotericin B (AmphoB; GoldBio) media to evaluate tolerance to an antifungal not used during the evolution experiment, though the ancestor was not included in this assay due to microplate capacity constraints. Doubling time (DT) and carrying capacity (*K*), used as proxies for population fitness, were estimated by fitting logistic growth models to OD_600_ time series using the “growthcurver” package v0.3.1 (Sprouffske and Wagner 2016). Both DT (inverse of growth rate *r*) and *K* are key life-history traits that characterize density-dependent population growth and have been used as fitness proxies in microbial evolutionary studies, with their relative contributions to fitness often depending on the environment (Sprouffske and Wagner 2016; Wei and Zhang 2019). Log-transformed DT and *K* were analyzed using linear mixed-effects models fitted with the “lme4” package v1.1.37 (Bates et al. 2015) with evolved population group, test medium, and their interaction as fixed effects and replicate as a random effect. Pairwise contrasts were derived from estimated marginal means using “emmeans”. More details on assay design, growth curve fitting, and modeling are provided in the **Supplementary Methods**.

## Acknowledgements

The authors thank the core facilities at the Center for Quantitative Life Sciences at Oregon State University for use of their computational and sequencing resources. We also thank members of the Burke Lab for feedback on earlier versions of the manuscript.

## Author Contributions

M.S.P. and M.K.B. designed the research; M.S.P., H.D., and E.S. performed the experiments; M.S.P., G.C., and M.K.B. analyzed genomic data; M.S.P. and M.K.B wrote the manuscript with input from all authors.

## Data availability

Sequence data for the 4S ancestor were previously published (Burke et al. 2020) and are available at NCBI SRA under BioProject PRJNA678990. Raw sequence data for all evolved populations generated in this study are available at NCBI SRA under BioProject PRJNA1544638. All R scripts used for data processing, statistical modeling, annotation, enrichment analysis, and figure generation, along with some input and output files needed to reproduce the analyses, are available on GitHub at https://github.com/MSandoval-Powers/Experimental-Evolution-Antifungal-Drug-Study. Larger data files, including raw and filtered SNP tables, are available on FigShare (DOI available upon publication).

