## Supplementary Methods and Figures S1-S7 for "Adaptive responses to antifungal selection depend on dose and drug combination in experimentally evolved yeast"

**Affiliations:**
[1]Department of Integrative Biology, Oregon State University,
2403 Cordley Hall,
2701 SW Campus Way,
Corvallis, OR 97331, USA

[*]**Authors for Correspondence:**
Megan Sandoval-Powers, Department of Integrative Biology, Oregon State University, Corvallis, Oregon, USA,
Molly K. Burke, Department of Integrative Biology, Oregon State University, Corvallis, Oregon, USA,

This file contains:
**Supplementary Methods**
**Supplementary Figures S1-S7**

Other supplemental files:
Tables S1-S7 are uploaded as a separate excel document.

### Supplementary Methods

#### Antifungal dose-response assays

To determine suitable doses for the evolution experiment, dose-response assays were conducted under conditions that mirrored the experimental evolution regime: 24-well plates, 1 mL culture volume, 30°C incubation with shaking. A range of concentrations of CAS (0–0.07 µM), CLO (0–3 µM), and their combinations were tested for effects on growth of the 4S ancestral population. To perform these assays, 1 mL of frozen 4S ancestor stock was spread onto yeast extract-peptone-dextrose (YPD) agar plates (10 g/L yeast extract, 20 g/L peptone, 20 g/L dextrose, 20 g/L agar) and incubated at 30°C for 48 hr. Cells were collected across the culture lawn using a sterile wooden applicator to capture population diversity and used to initiate three independent overnight cultures (biological replicates) in 10 mL YPD broth. Cultures were grown overnight at 30°C with shaking at 200 rpm. Each overnight culture was standardized to an optical density at 600 nm ($OD_{600}$) of ~ 0.1 in YPD medium containing the desired concentrations of CAS, CLO, CASCLO, or 0.05% DMSO as a control, resulting in three biological replicates per treatment condition. The DMSO control was included because all antifungal drugs were dissolved in DMSO, and 0.05% reflects the final solvent concentration present in the highest drug-containing wells, ensuring that any effects of DMSO itself were accounted for. A 1 mL volume of each standardized culture was added into sterile, flat-bottom 24-well plates (Corning, Corning, NY, USA; Cat. No. 09-761-146), and plates were incubated at 30°C with shaking at 200 rpm. Growth was quantified by measuring $OD_{600}$ at 24 hr and 48 hr using an Eppendorf BioSpectrometer (Eppendorf, Hamburg, Germany) and expressed as mean relative $OD_{600}$ (% of control) ± SD of the three biological replicates.

#### Yeast base population

The ancestral *S. cerevisiae* 4S population used in this study originated from an outcrossed, highly recombined base population constructed by crossing four diverse haploid founder strains (YPS128, Y12, DBVPG6765, and DBVPG6044) from the SGRP collection (Cubillos et al. 2009). To facilitate efficient mating and diploid recovery, these founder strains were genetically engineered as described by Linder et al. (2020) (strains donated by Anthony Long, UC Irvine, CA, USA). Briefly, *MATa* strains carried *ycr043C::NatMX*, *ura3::KanMX-barcode*, *ho*Δ, whereas *MATα* strains carried *ycr043C::HygMX*, *ura3::KanMX-barcode*, *ho*Δ, allowing selection of mating products while preventing mating-type switching. After initial crossing, the 4S base population was carried through an additional 12 sexual cycles to increase recombination, expand genetic diversity, and promote adaptation to laboratory conditions (Burke et al. 2020; Phillips et al. 2021). This well-characterized, genetically diverse base population served as the ancestor for all experimental populations in the current study.

#### Whole-genome sequencing library preparation

Genomic DNA (gDNA) was extracted from all samples designated for sequencing, diluted to 6–9 ng/µL, and stored at –20°C in 96-well PCR plates. Tagmentation was performed using the Illumina Tagment DNA Enzyme and Buffer Kit (Illumina, San Diego, CA, USA; Cat. No. 20034197). A tagmentation master mix was prepared by combining TD buffer and TDE enzyme in proportions scaled to the number of samples with additional volume for pipetting. Six microliters of the master mix were dispensed into new PCR plates, followed by 4 µL of diluted gDNA per well using a multichannel pipette. Plates were sealed, mixed by pipetting, briefly centrifuged, and incubated at 55°C for 5 minutes in a thermal cycler.

Following tagmentation, the DNA fragments were purified using MagBio HighPrep PCR magnetic beads (MagBio Genomics, Gaithersburg, MD, USA; Cat. No. AC-60050) to remove enzymes, buffers, and other reaction components prior to PCR amplification. Beads were equilibrated to room temperature prior to use and added at 1.8 times the sample volume of tagmented DNA and mixed thoroughly. Following a 5-minute room temperature incubation, plates were placed on a magnetic rack to clear, and the supernatant was carefully removed. Two washes with 180 µL fresh 80% ethanol per well were performed, each followed by brief incubation on the magnet and ethanol removal. DNA was eluted with 25 µL Qiagen Buffer EB (Qiagen, Hilden, Germany; Cat. No. 19086) per well and transferred to fresh PCR plates for further processing.

After purification, DNA was labeled with custom dual-index primers from a Nextera Index Kit (Illumina; Cat. No. FC-121-1011) and amplified by PCR using a KAPA HiFi HotStart Library Amplification Kit (Roche, Wilmington, MA, USA; Cat. No. KK2612) with custom dual-index primers (i5 and i7) from the Nextera Index system (Illumina; Cat. No. FC-121-1011). The i5 (forward) and i7 (reverse) primers appended Illumina adapter sequences and unique dual indices (barcodes) to each library, enabling multiplexed sequencing and subsequent demultiplexing of reads. Each 20 µL reaction contained 2 µL of Nextera PCR primer mix (10 µM each, 10 µL of 2X KAPA MasterMix, 2 µL of i5 index primer (0.5 µM), 2 µL of i7 index primer (0.5 µM), and 4 µL tagmented DNA. Reactions were mixed gently by pipetting, briefly centrifuged, and amplified using a thermal cycler with an initial extension at 72°C for 3 minutes, denaturation at 98°C for 30 seconds, followed by 12 cycles of 98°C for 10 seconds, 63°C for 30 seconds, and 72°C for 3 minutes.

Amplified libraries were quantified via a Qubit fluorometer (Thermo Fisher Scientific, Waltham, MA, USA) and samples were pooled based on concentration data to normalize input. Pooled libraries were further purified using a Qiagen MinElute PCR Purification Kit (Cat. No. 28004) and size selected via 2.5% agarose gel electrophoresis. DNA fragments between ~450–600 bp were excised and extracted using the Qiagen MinElute Gel Extraction Kit (Cat. No. 28604). Purified libraries were quantified again with Qubit, diluted, and sequenced on PE150 lanes in an Illumina NextSeq 2000 at the Center for Quantitative Life Sciences (Oregon State University, Corvallis, OR.)

**Assessment of variation prior to modeling**

PCA was performed in R v4.5.1 (R Core Team 2025) using the ‘prcomp’ function on allele frequencies standardized to unit variance across all polymorphic sites. To examine treatment-specific divergence, separate PCAs were also performed on subsets of populations corresponding to each experimental contrast (e.g., Low CAS vs. High CAS populations), with populations annotated by treatment and timepoint. SFS were calculated for each population at the final timepoint ($T_{14}$) using allele frequencies at all filtered polymorphic SNPs.

For the chromosome VIII analysis, read depth at each filtered SNP position was normalized to the genome-wide median coverage of that sample to control for differences in library depth. For each replicate population at each timepoint, the median relative coverage across chromosome VIII was summarized. Median coverage in 20 kb window along the chromosome were also calculated to show whether coverage rose along the whole chromosome or only in segments. Windows below 20 kb and above 540 kb were dropped, because repetitive sequence near the chromosome ends inflated coverage in every sample, including Controls. Since pooled sequencing data reflects an average across many cells, these patterns were treated as indirect evidence of copy number change.

**Growth assay protocol**

Low CAS, High CAS, and ancestral populations were assayed in control, Low CAS, and High CAS media; Low CLO, High CLO, and ancestral populations in control, Low CLO, and High CLO media; and CASCLO and ancestral populations in control, High CAS, and CASCLO media. High CAS and CASCLO populations were additionally assayed in control, High CAS, CASCLO, and 2 μM Amphotericin B (AmphoB) media to assess cross-tolerance. Each assay included all 12 evolved replicate populations per group and, where applicable, three biological replicates of the ancestral population.

Populations were revived from 600 μL of frozen stocks on YPD agar, and cells were collected across culture lawns to initiate overnight liquid cultures in 10 mL YPD at 30°C with 200 rpm shaking. Overnight cultures were diluted to an $OD_{600}$ of ~ 0.1 in the appropriate drug or control medium and transferred to individual wells of a 96-well flat-bottom microplate (Corning; Cat. No. 351172) in 200 μL volumes. Plates were covered with a lid and incubated at 30°C in a microplate reader (Tecan Spark) and $OD_{600}$ was measured every 30 minutes over 48-hr, with orbital shaking before each reading to minimize cell settling. Measurements were taken at four positions within each well and averaged prior to analysis.

Wells in which populations failed to grow or reach stationary phase, particularly ancestral populations in CAS and CASCLO media, were excluded from growth curve fitting because the logistic model could not reliably estimate DT and $K$; failure to grow was noted separately and is reported as a qualitative outcome. To directly compare populations evolved under the same drug at different concentrations (e.g., Low CAS vs. High CAS), data from the respective assays were combined and analyzed using the same model structure described in the main text. Assay was tested for significance and included in the model when appropriate to account for potential batch effects, and pairwise contrasts were Tukey-corrected for multiple comparisons.

#### Supplementary Figures

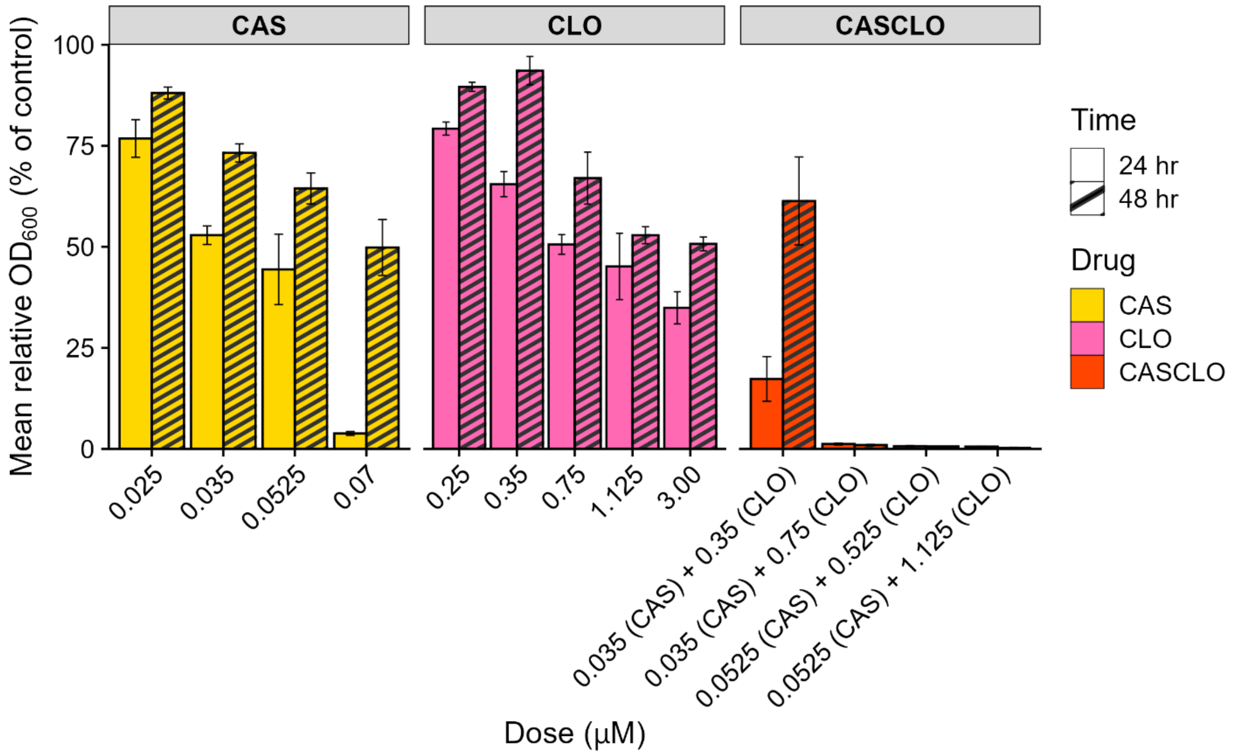


**Figure S1. Dose-response assays identify low and high antifungal concentrations used for experimental evolution.** Mean relative growth of the ancestral 4S population, measured as $OD_{600}$ and expressed as a percentage of the 0.05% DMSO control, is shown at 24 hr (solid bars) and 48 hr (striped bars) across the range of drug concentrations tested. Responses to caspofungin acetate (CAS) are shown as yellow bars, clotrimazole (CLO) as pink bars, and the combination (CASCLO) as orange bars. Error bars represent ± standard deviation of three biological replicates. The doses selected for the evolution experiments were 0.035 µM and 0.07 µM for CAS, 0.35 µM and 3.0 µM for CLO, and 0.035 µM CAS + 0.35 µM CLO for the combination treatment.

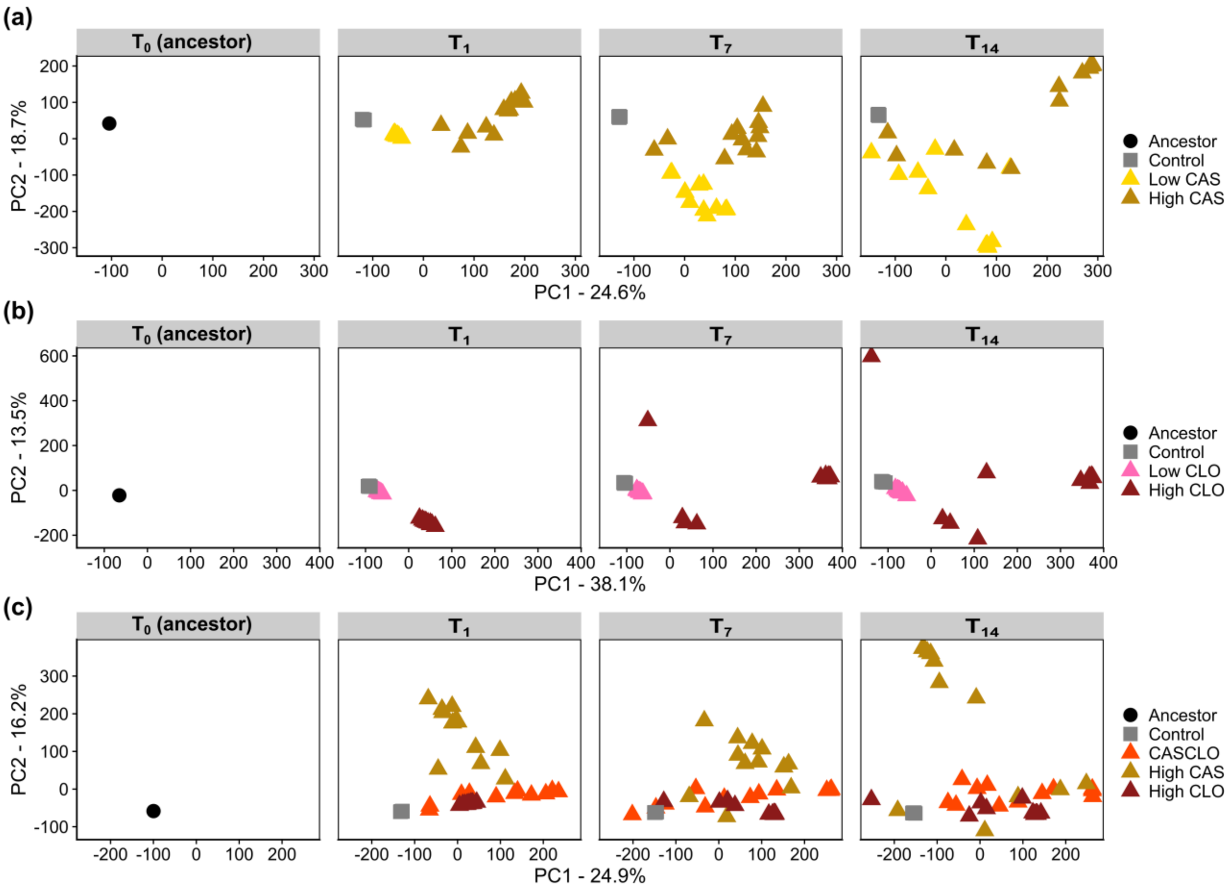


**Figure S2. Principal component analysis of genome-wide allele frequencies reveals treatment-specific divergence over time.** Principal component analysis was performed on allele frequencies across all polymorphic sites (*n* = 64,764) for each contrast, with populations plotted by timepoint ($T_0$, $T_1$, $T_7$, $T_{14}$), where $T_0$ represents the ancestral population. Separate analyses were performed for each contrast: (a) the CAS intensity comparison (Low CAS, High CAS, Control), (b) the CLO intensity comparison (Low CLO, High CLO, Control), and (c) the single- versus combined-drug comparison (CASCLO, High CAS, High CLO, Control). Each point represents one replicate population, colored by treatment. Axis labels give the percent of total variance explained by PC1 and PC2.

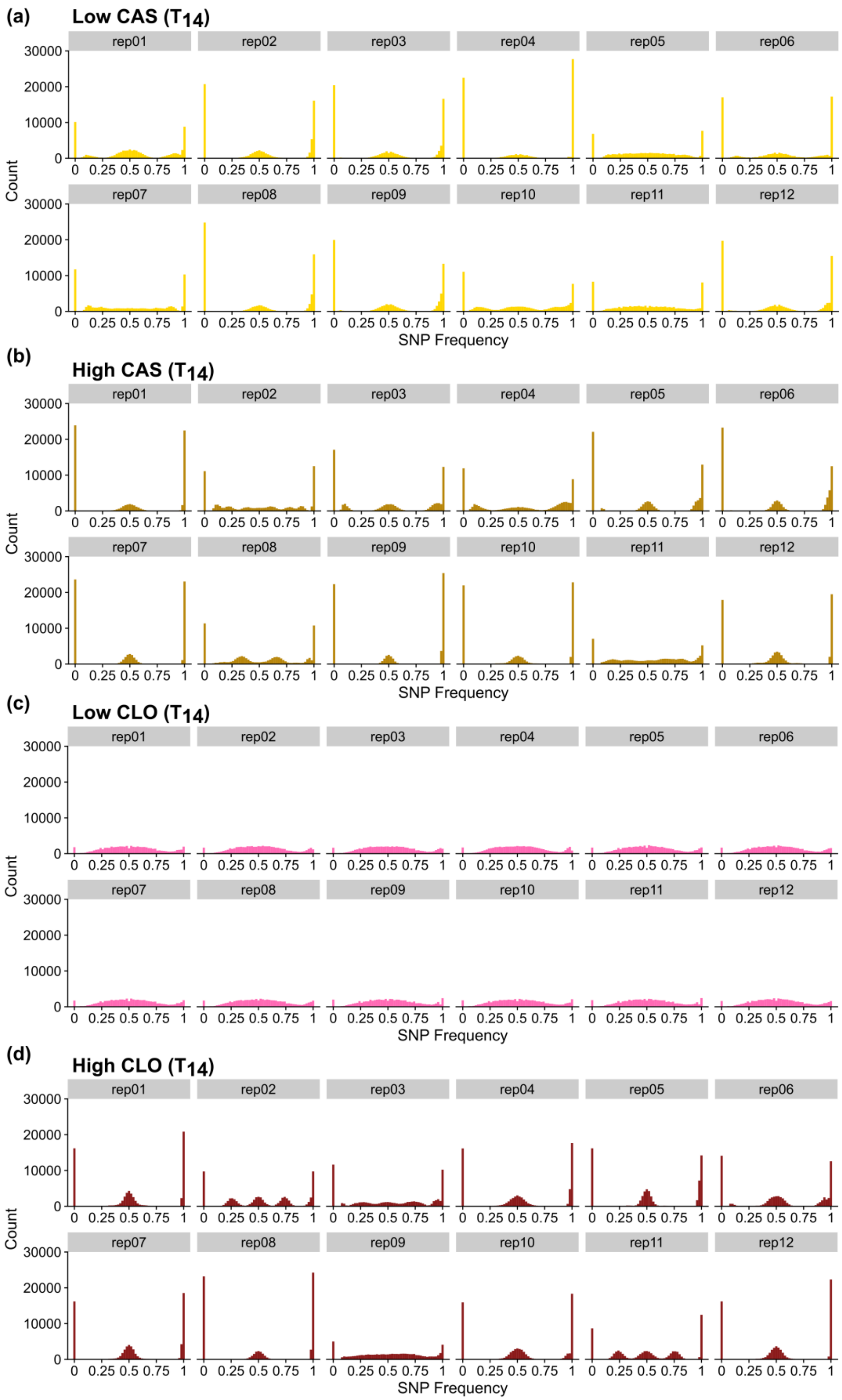
(a)
Low CAS (T14)
rep01
rep02
rep03
rep04
rep05
rep06
rep07
rep08
rep09
rep10
rep11
rep12
Count
SNP Frequency
(b)
High CAS (T14)
(c)
Low CLO (T14)
(d)
High CLO (T14)

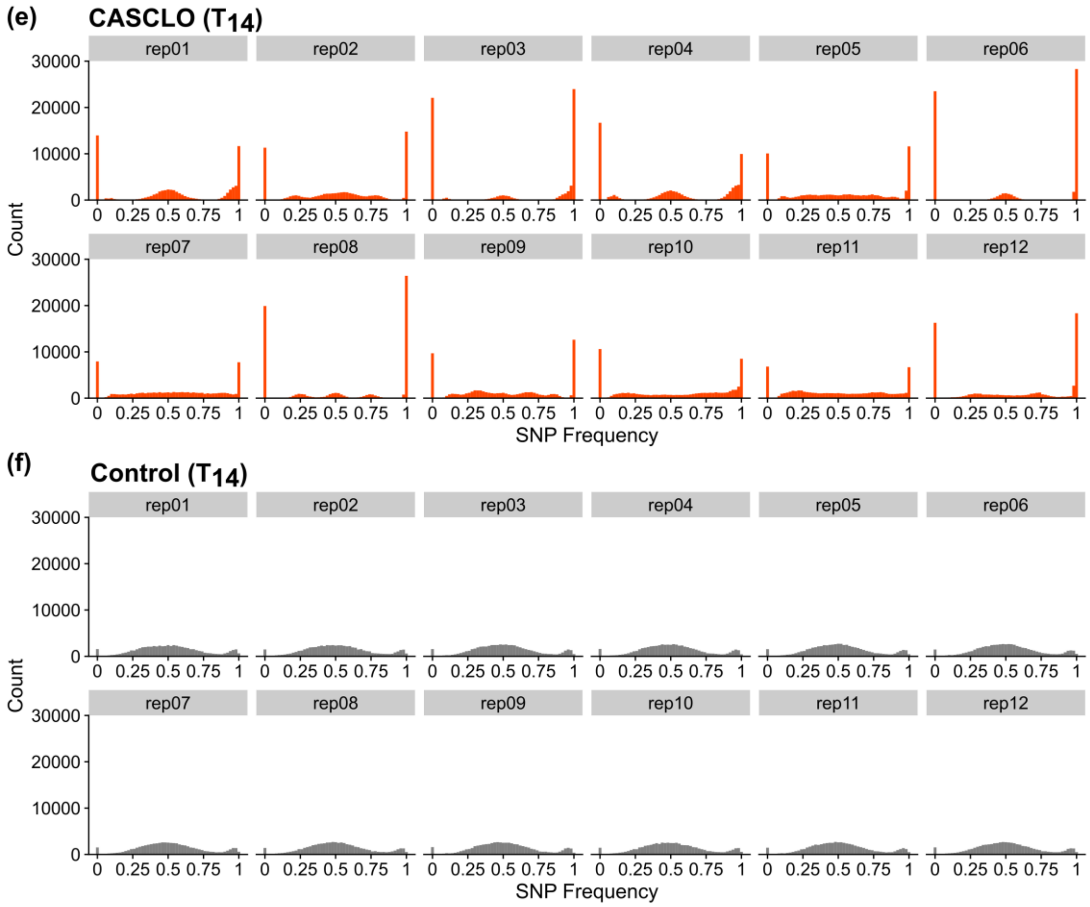


**Figure S3. Allele frequency distributions across replicate evolved populations.** Site frequency spectra (SFS) were calculated from filtered polymorphic SNPs ($n$ = 64,764) at the final timepoint ($T_{14}$) to characterize genome-wide allele frequency distributions within evolved populations. Histograms show the number of SNPs observed across allele frequency bins ranging from 0 to 1, with each facet representing an individual replicate population. Panels show populations evolved under Low CAS (a), High CAS (b), Low CLO (c), High CLO (d), CASCLO (e), and Control populations (f).

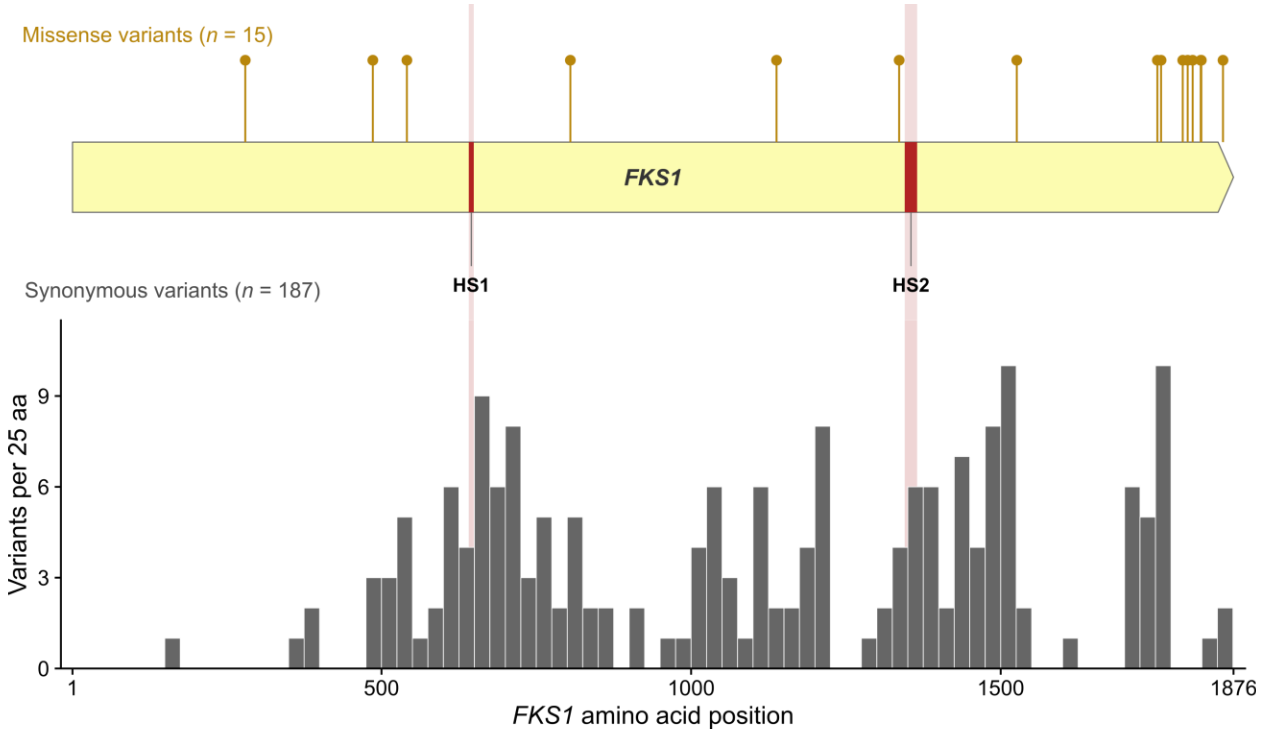


**Figure S4. *FKS1* candidate variants in CAS populations were distributed across the gene rather than concentrated in known resistance hotspots.** Positions of missense (top; *n* = 15) and synonymous (bottom; *n* = 187) candidate variants along the FKS1 protein (amino acids 1–1876). Synonymous variants are shown as counts per 25 amino acid bin. Shaded regions mark hotspot regions HS1 (residues 641–649) and HS2 (residues 1345–1365), where most known echinocandin resistance mutations occur (Perlin 2007; Johnson et al. 2011). No candidate missense variants fell within either hotspot.

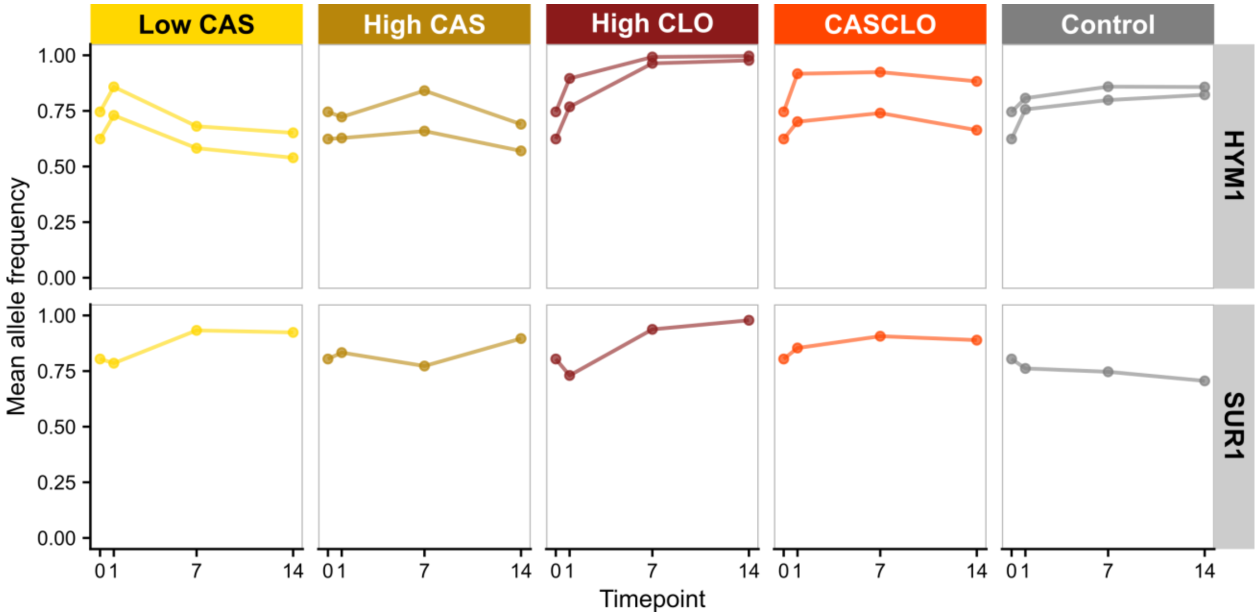


**Figure S5. Allele frequencies at *HYM1* and *SUR1* candidate missense variants increased most strongly under High CLO selection.** Allele frequency trajectories at the candidate missense variants in *HYM1* (2 variants, top row) and *SUR1* (1 variant, bottom row) across five treatments: Low CAS, High CAS, High CLO, the CASCLO combination, and the no-drug Control. Each line is one variant, plotted as the mean allele frequency across the 12 replicate populations per treatment. $T_0$ is the ancestral population value and is identical across panels. Low CLO is not shown as it was omitted from gene-level analyses.

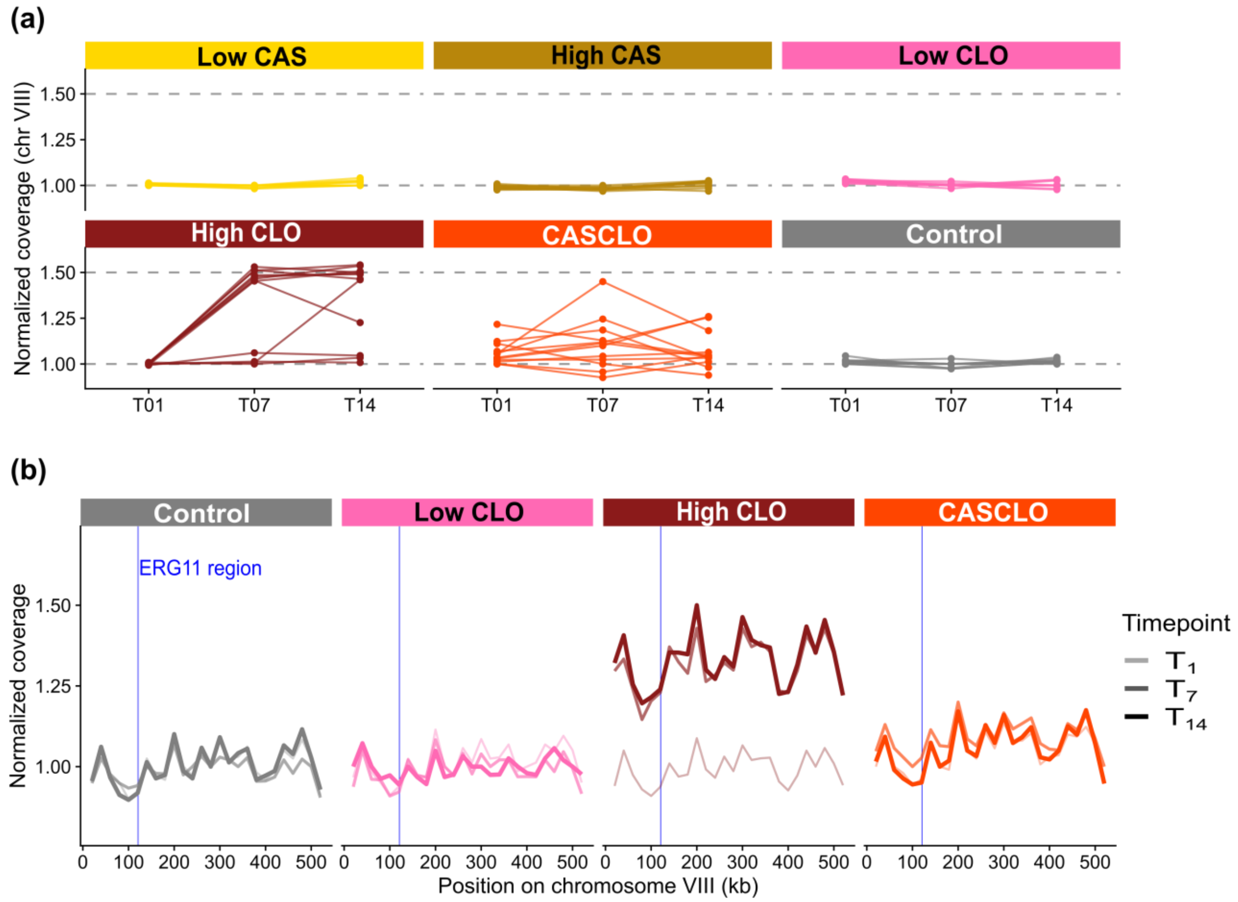


**Figure S6. Increased chromosome VIII coverage under High CLO selection.** (a) Normalized coverage of chromosome VIII for each replicate population across timepoints. Coverage at each SNP position was divided by the genome-wide median coverage of that sample, such that 1.0 corresponds to ancestral copy number and 1.5 to one additional chromosome copy in a diploid (dashed lines). (b) Normalized coverage along chromosome VIII in 20 kb windows, shown as the median across replicates within each treatment. Line opacity and weight indicate timepoint. Vertical line marks the position of *ERG11* (chromosome VIII: 120,091–121,683). Positions below 20 kb and above 540 kb were excluded because repetitive subtelomeric sequence inflated coverage in all samples, including Controls.

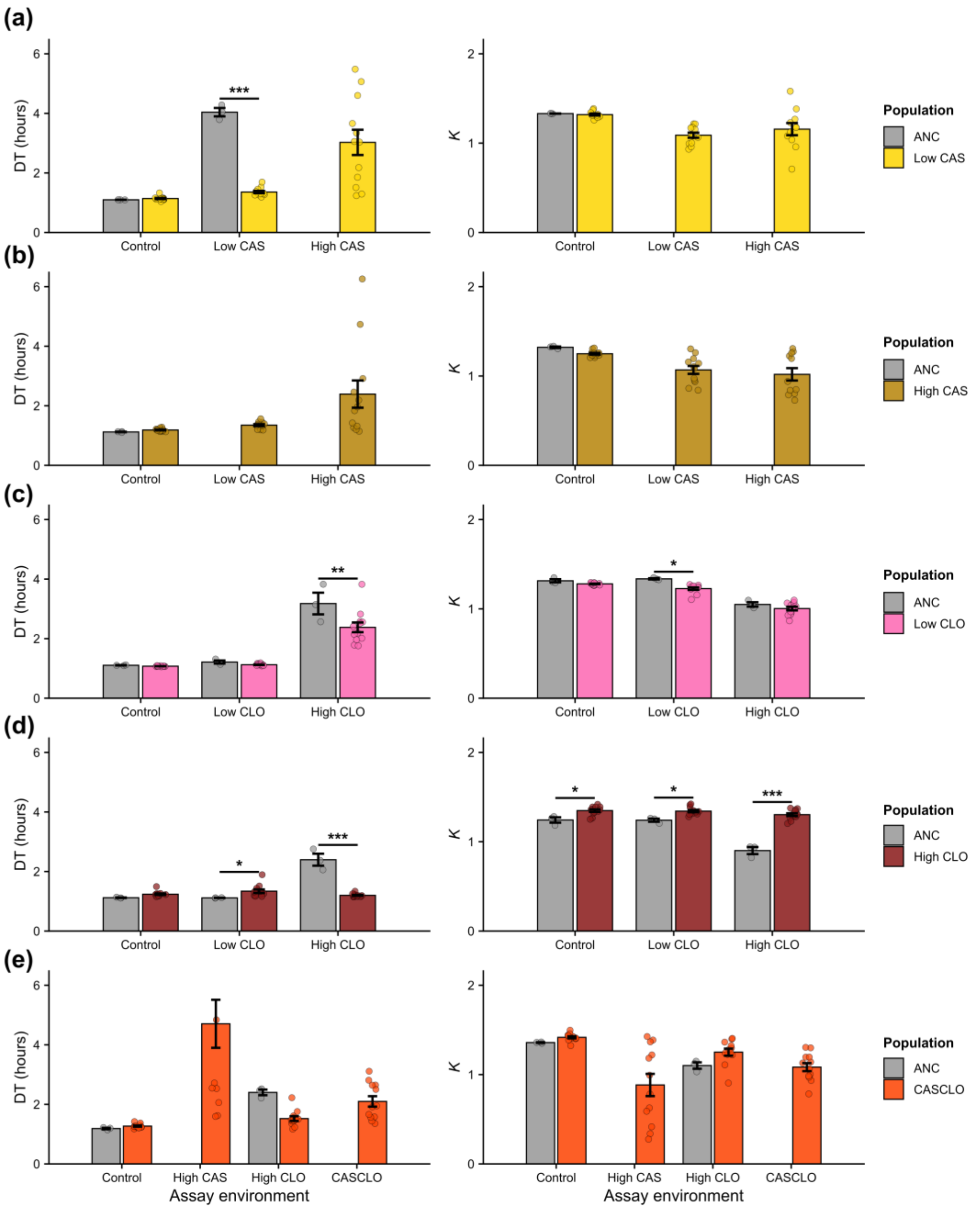


**Figure S7. Evolved populations grow faster than the ancestor under high- and combined-drug conditions.** Doubling time (DT) and carrying capacity (*K*) were measured from 48 hr growth assays of evolved populations from the final timepoint ($T_{14}$) and the ancestor (ANC) across control and drug-containing media. DT is shown in the left column and *K* in the right column for Low CAS (a), High CAS (b), Low CLO (c), High CLO (d), and CASCLO (e) populations. Within each panel the x-axis gives the assay environment, and grey and colored bars show the DT and *K* values for ancestor and the evolved populations, respectively. Values are shown as mean ± SE across replicate populations. In environments where the ancestor failed to grow, grey bars are omitted because DT or *K* could not be reliably estimated. Statistical significance was determined using linear mixed-effects models applied to log-transformed DT and *K* values with replicate included as a random effect, and pairwise contrasts were Tukey-corrected. Asterisks indicate significant pairwise differences between the ancestor and evolved populations within each assay environment (*$p < 0.05$; **$p < 0.01$; ***$p < 0.001$).
